# RNA Splicing Junction Landscape Reveals Abundant Tumor-Specific Transcripts in Human Cancer

**DOI:** 10.1101/2024.02.06.579246

**Authors:** Qin Li, Ziteng Li, Bing Chen, Jingjing Zhao, Hongwu Yu, Jia Hu, Hongyan Lai, Hena Zhang, Yan Li, Zhiqiang Meng, Zhixiang Hu, Shenglin Huang

## Abstract

RNA splicing is a critical process governing gene expression and transcriptomic diversity. Despite its importance, a detailed examination of transcript variation at the splicing junction level remains scarce. Here, we perform a thorough analysis of RNA splicing junctions in 34,775 samples across multiple sample types. We identified 29,051 tumor-specific transcripts (TSTs) in pan-cancer, with a majority of these TSTs being unannotated. Our findings show that TSTs are positively correlated with tumor stemness and linked to unfavorable outcomes in cancer patients. Additionally, TSTs display mutual exclusivity with somatic mutations and are overrepresented in transposable element-derived transcripts possessing oncogenic functions. Importantly, TSTs can generate neoepitopes that bind to MHC class I molecules for immunotherapy. Moreover, TSTs can be detected in blood extracellular vesicles from cancer patients. Our results shed light on the intricacies of RNA splicing and offer promising avenues for cancer diagnosis and therapy.

**In brief:** This study thoroughly analyzed RNA splicing junctions in 34,775 samples and identified 29,051 tumor-specific transcripts (TSTs), which may serve as novel cancer driver genes, neoantigens, and circulating biomarkers.

**Graphical abstract:** 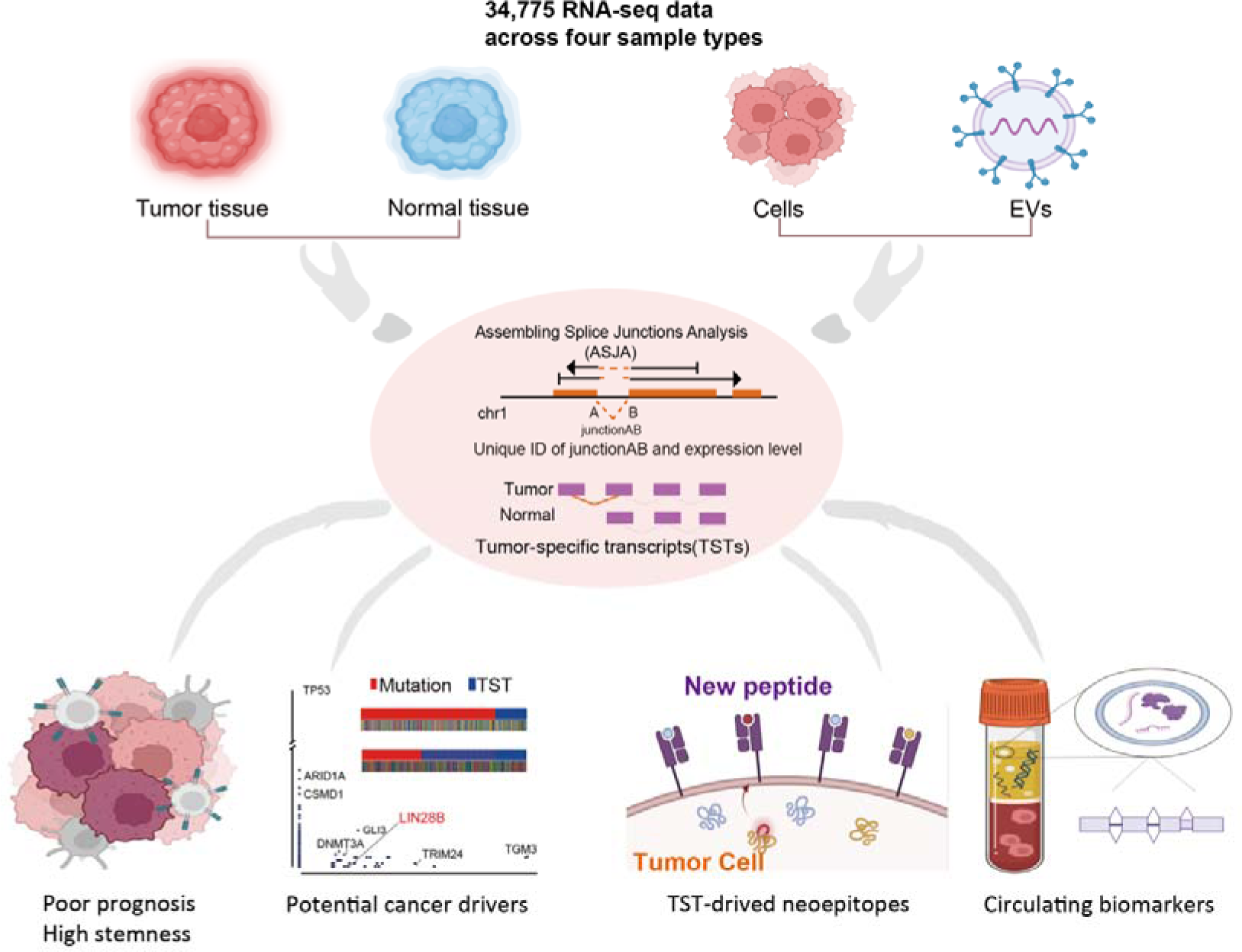

## Introduction

RNA molecules play a pivotal role in the transmission and regulation of genetic information. Recent advancements in deep RNA sequencing technologies (RNA-Seq) and bioinformatics analysis have unveiled the intricate and multifaceted nature of the human transcriptome^1,2^. RNA splicing, a crucial enzymatic process in eukaryotic cells, removes segments of premature RNA to produce mature mRNA, and is a key mediator of transcript diversity and gene expression regulation^3^. The emergence of RNA-Seq technology has revolutionized transcriptome analysis by providing unparalleled resolution and scope^4^. However, a systematic and precise analysis of transcript variation at the splicing junction level remains insufficient for comprehensive understanding.

Dysregulation of RNA splicing is a critical mechanism in tumor pathogenesis^5^. Recent analyses of genetic alterations in cancer and their relationship to the transcriptome and epigenome have revealed numerous ways in which splicing is altered in cancer cells^6-8^. Efforts to compare the transcriptomes of tumor and normal tissue, such as The Cancer Genome Atlas, demonstrate that many cancers exhibit aberrant splicing, including changes in usage of annotated transcript isoforms and an increased use of aberrant unannotated splicing events^9^. Existing studies have focused on specific types of splicing events associated with abnormal transcripts, such as 3’ UTR splicing driving tumorigenesis^10^ and exonic splicing analysis revealing tumor-specific events^11^. Our previous works have identified hundreds of tumor-specific transcripts (TSTs) in patients with hepatocellular carcinoma^12^ and triple-negative breast cancer^13^, and characterized the functional roles of LIN28B-TST^14^ and MARCO-TST^13^. We anticipate that various forms of TSTs with aberrant splicing due to epigenetic abnormalities or post-transcriptional processing may commonly exist in cancer. However, widespread identification and exploration of their potential functions remain incomplete.

In this study, we systematically detected RNA splicing junctions in 34,775 samples from five different sample types using an optimized RNA splicing junction detection tool. We identified 870,389 splicing events, with nearly half of these splicing junctions being unannotated. We further discovered 29,051 tumor-specific transcripts (TSTs) from 32,848 splicing sites that are specific to tumors. Our findings suggest that TSTs could serve as potential drivers of tumorigenesis and sources of novel tumor antigens. Moreover, we performed extracellular vesicles (EVs) RNA-Seq and explored the splicing sites in EVs for tracking tissue sources and cancer liquid biomarkers.

## Results

### Identification of RNA splicing junctions across different human tissues and cells

To construct the landscape of splicing events across different human sample types, we collected RNA-Seq data of 17,382 normal samples (30 tissue types), 9,581 cancer samples and 702 matched normal samples (33 tissue types), 1128 extracellular vesicle (EV) samples (4 biofluids), 1046 cancer cell lines and 4936 single cells (6 cell types) (Supplementary Table 1). We developed an optimized RNA splicing junction detection tool (Assembling Splice Junctions Analysis, ASJA) to analyze the RNA-seq data (Fig. 1a). ASJA identified unique splicing junction position in all regions of the genome and quantified junction expression based on assembling transcripts allowing comparison among samples. In total, we identified 870,389 splicing events including 333,349 in normal tissues, 667,215 in cancer tissues, 472,193 in EVs and 526,982 in cells (Fig. 1b). There are 246,652 splicing events present across all the sample types (Fig. 1b).

**Figure 1.**
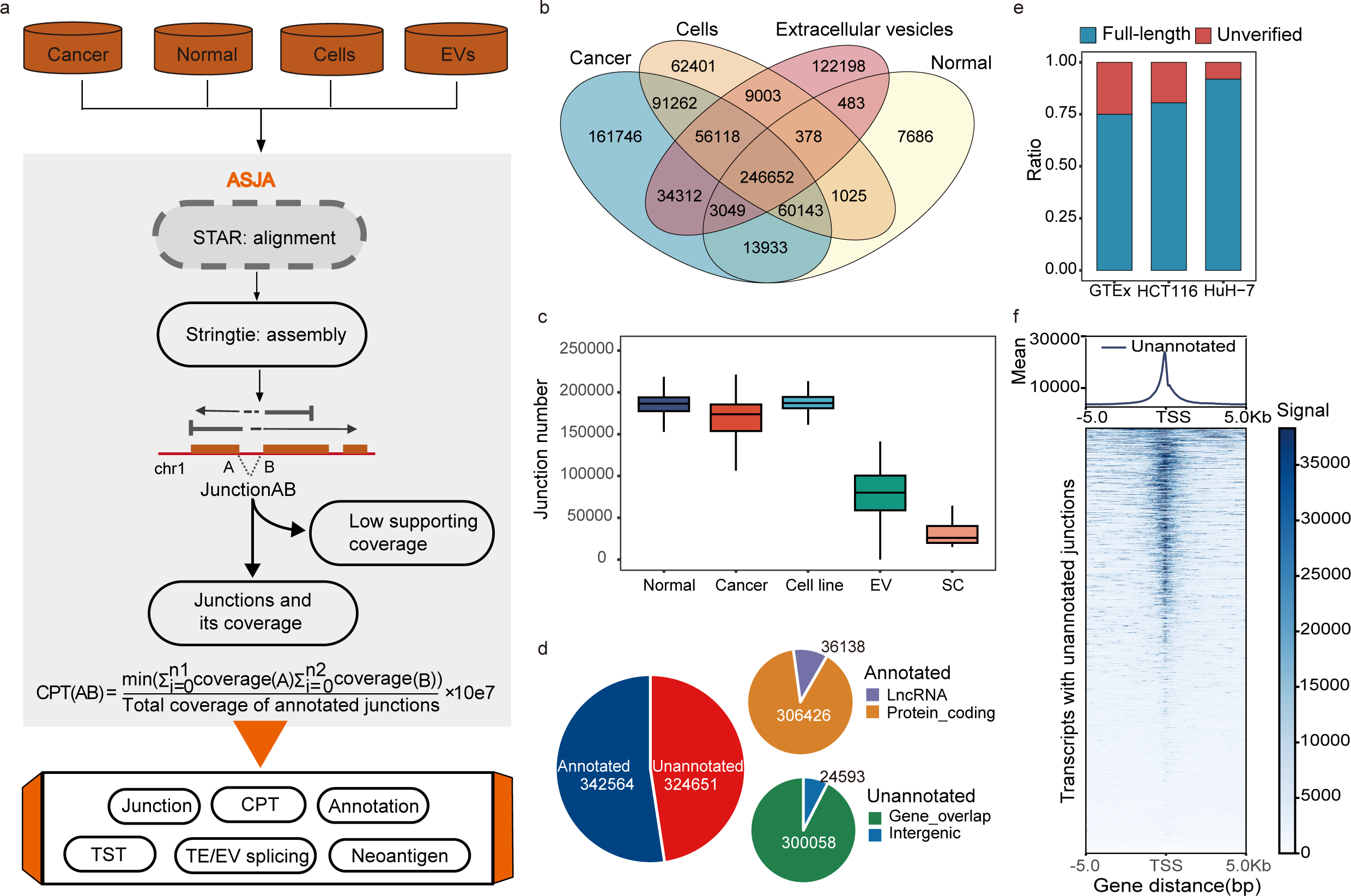
Identification of RNA splicing junctions across different human tissues and cells. a, Schematic diagram of the workflow from sample collection to splicing sites detection. b, The number and types of splicing sites identified in different types of samples. “Cells” include 1046 cancer cell lines and 4936 single cells here. c, Number of splicing sites detected in each sample of six sample types. Each dot represents a sample. The horizontal line in the box plot indicates the median number of linear splicing sites detected in each sample type. d, Category of splicing sites. The category of annotated and unannotated splicing sites are shown in two additional pie charts. e, Proportion of splicing sites detected by second-generation RNA-seq data validated by third-generation sequencing. f, ATAC signal enrichment near transcription start sites of transcripts containing unannotated splicing junctions. Rows represent chromatin openness and are sorted by ATAC signal enrichment values. Color indicates normalized signal enrichment levels. Abbreviation: CPT, coverage per ten million; TST, tumor-specific transcripts. TE/EV splicing, splicing derived from transposable elments/extracellular vesicles. SC, single cells.

For each sample type, we obtained an average of 186,567 splicing events per normal tissue, 181,711 per adjacent normal tissue, 173,747 per cancer tissue, 80,109 per EV sample, 189,171 per cell line and 25,983 per single cell (Fig. 1c). Generally, splicing junctions detected in normal tissues were more numerous compared to cancer. Notably, about half of the splicing events (324,651, 48.65%) detected in cancer tissues are unannotated (Fig. 1d). Among annotated junctions, 306,426 (89.45%) were from protein-coding genes, and only 36,138 were from lncRNA (Fig. 1d). Among unannotated junctions, 300,058 (92.42%) were overlapped with gene which may represent new isoforms for host genes, and 24,593 were from intergenic regions (Fig. 1d).

To demonstrate the robustness of splicing events detected using ASJA from the short-read RNA-seq data, we made the comparison with long-read RNA-seq data. About 75% of the junctions could be observed in the long-read RNA-seq data from GTEx project. We also performed the short-read and direct full-length RNA-seq of HuH-7 and HCT 116 cells. For the detected junctions from short-read RNA-seq, about 80% in HCT 116 and 90% in HuH-7 could be validated from direct RNA-seq data (Fig. 1e). Moreover, the analysis of chromatin accessibility profiles of 23 types of primary human cancers, represented by 410 tumor samples derived from 404 donors from TCGA, revealed that clear ATAC-seq peaks were detected around the transcriptional start site of parental transcripts with unannotated junctions (Fig. 1f). Those results suggested that the identified RNA splicing junctions were derived from bona fide transcripts.

### Characterization of RNA splicing junctions

To identify splicing type for each junction, alternative splicing events from those junctions were extracted. We identified a total of 114,490 alternative splicing events, of which alternative promoter (AP) events accounted for the majority (45,955, 40.14%) (Fig. 2a). Notably, a substantial number of splicing events exit in the intronic and intergenic regions (Fig. 2b). Even in spliced exonic regions including coding sequences (CDS) and untranslated region (UTR, 3’ and 5’), there could still occur splicing events (Fig. 2b). Supporting this observation, some recent studies provided experimental evidences for the true existence and functional roles of atypical splicing sites^10,11^, suggesting these junctions are attractive potential tumor targets and worth further investigation.

**Figure 2.**
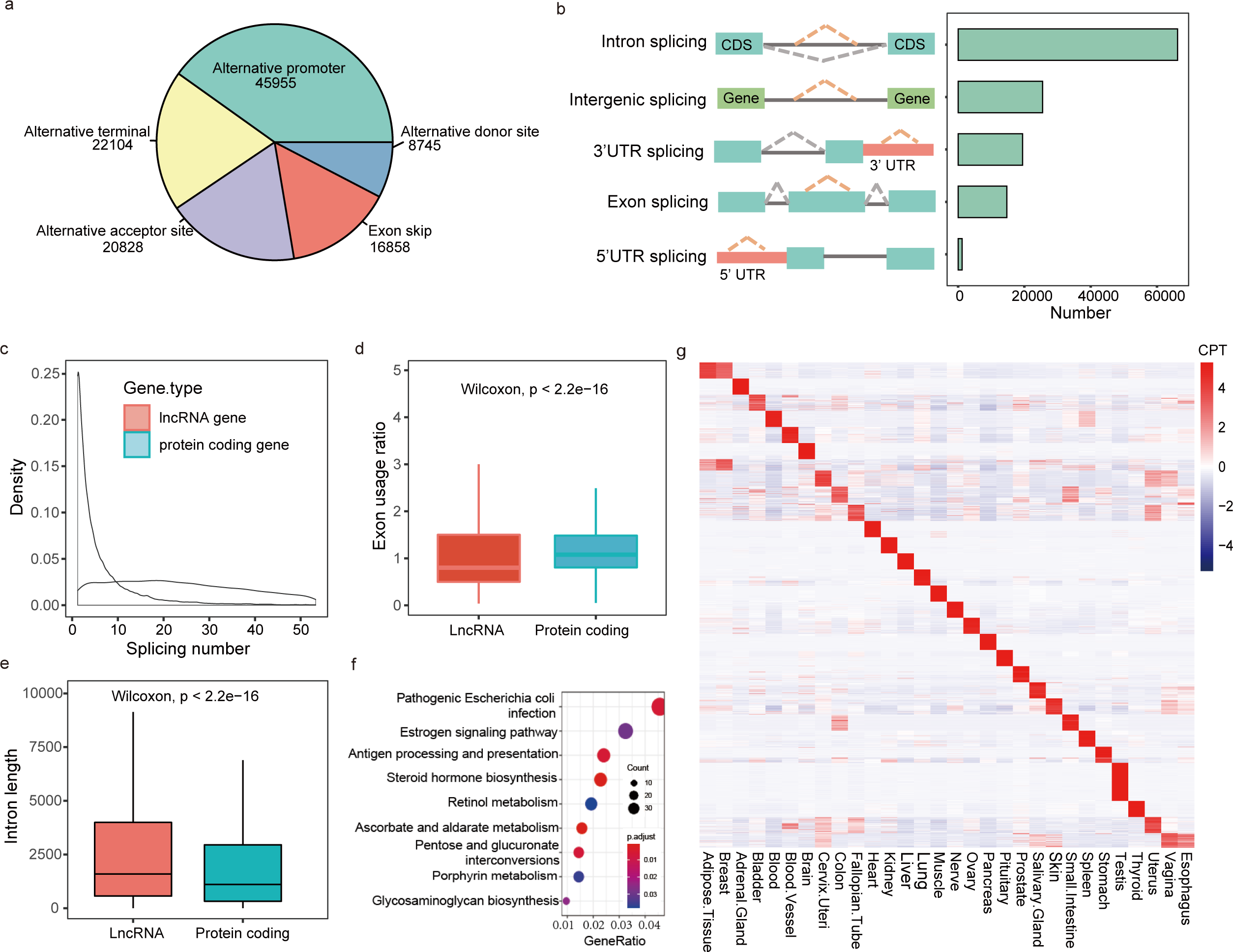
Characterization of RNA splicing junctions. a, Types and numbers of alternative splicing sites. b, Types and numbers of atypical splicing sites. Atypical splicing refers to the junctions that both donor and acceptor sites are located in exons, introns or intergenic regions. c, Density plot of splicing site numbers for protein-coding genes and lncRNA genes. d, Comparison of exon usage rates between protein-coding genes and lncRNA genes (Wilcoxon test, p < 2.2×10−16). Exon usage rate = gene splicing number / exon number. e, Comparison of intron lengths between protein-coding genes and lncRNA genes (Wilcoxon test, p < 2.2×10−16). f, Enriched KEGG pathways of frequently TST-derived genes (exon usage rate > 2). The exon usage rate for each gene was calculated as the number of splicing junctions divided by the number of exons. g, Heatmap shows the top 100 splicing sites with highest tissue-specificity scores. The expression median of each tissue was normalized for visualization purposes.

We further investigate splicing characteristics for different gene types and found that protein-coding genes had many more splicing events than lncRNA genes (median: 24 vs median: 3) (Fig. 2c). Even after excluding exon number, protein-coding genes still had higher exon usage rates than lncRNA genes (Wilcox test, p < 2.2×10−16) (Fig. 2d). Notably, intronic lengths of lncRNA genes were observed to be longer than those of protein-coding genes (Fig. 2e), suggesting that intronic length may affect splicing. Among these spliced genes, a group of genes were spliced frequently (exon usage ratio>2). These genes were found to be involved in different biosynthesis and metabolic pathways (Fig. 2f).

Additionally, we found 143,463 splicing events under tissue-specific splicing regulation across GTEx tissue types (Fig. 2g). We next asked whether the splicing events marked tissue-specific genes. Tissue-specific (TS) score was calculated for each junction and gene across 30 normal tissues (see Methods). We found 90,768 splicing events under strong tissue-specific regulation (TS score>0.95, Supplementary Table 2). By checking the tissue specificity of gene expression patterns of their parental genes, we found that tissue-specific splicing junctions were significantly enriched for tissue-specific genes compared to random expectation (p-value<2.2e-16, Fisher’s exact test). However, we observed that some genes did not show tissue-specific expression, but their splicing sites had obvious tissue preferences. For example, the splicing site chr22:37688360|37688850:+ from protein-coding gene *NOL12* was specifically expressed in testis (Extended Data Fig. 1a), while its parental gene did not show tissue specificity (Extended Data Fig. 1b).

### Discover tumor-specific transcripts (TSTs) in pan-cancer

Although the number of splicing junctions in each cancer tissue was less than that of normal tissue, a substantial number of splicing junctions exclusively exist in cancers, suggesting the presence of abundant tumor-specific transcripts (TSTs). We performed stringent criteria to identify the tumor-specific splicing events. Only those junctions frequently expressed (>5%) and specifically expressed or 10-fold higher in cancer compared to the maximum of adjacent and normal tissues are considered as tumor-specific junctions (TSJs) (Fig. 3a). In total, we discovered 32,848 TSJs derived from 29,051 TSTs (Supplementary Table 3). TSTs are frequently detected across 33 cancer types from 270 to 6842, of which ovary cancer (OV) and testicular germ cell tumors (TGCT) have the largest numbers of TSTs (Fig. 3b). Most of the TSTs (28,200, 97.07%) are un-annotated (Fig. 3c), and more than half of TSTs (16,284, 56.12%) were generated from intergenic region (Fig. 3d). The transcriptional start site of TST exhibits open chromatin features, implicating disrupted epigenetic regulation in tumorigenesis as a possible mechanism underlying aberrant TST expression (Fig. 3e). Moreover, most of these TSTs identified from cancer tissues were also present in cancer cell lines (25,808, 88.95%), suggesting that TSTs mainly from cancer cells. It is noteworthy that these TSTs included LIN28B-TST and MARCO-TST that we previously reported^13,14^.

**Figure 3.**
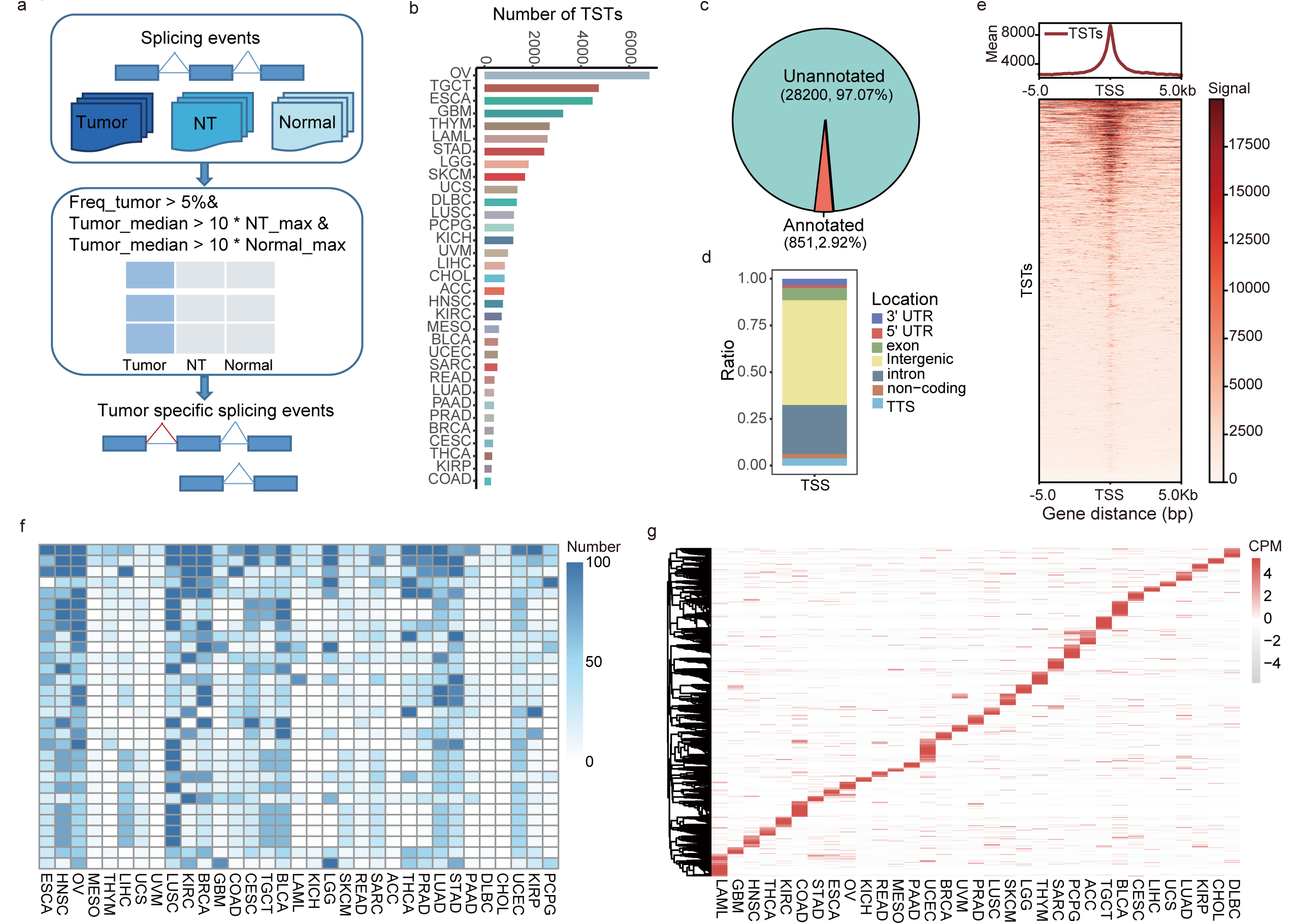
Discover tumor-specific transcripts (TSTs) in pan-cancer. a, Workflow for the identification of tumor-specific junctions (TSJs). Splicing junctions with absolute high expression in cancer samples were identified as TSJs, and only those with expression frequency greater than 5% were retained. b, Number of TSJs identified in each TCGA cancer type. c, Number and proportion of annotated and unannotated TSJs. d, Genomic distribution of donor and acceptor sites of TSJs. “Promoter-TSS” represents the region from -1kbp to +100bp near the transcription start site, and “TTS” represents the region from -100bp to +1kbp near the transcription termination site. e, ATAC-seq signal enrichment within 5kb of transcription start sites of TSTs. f. Expression of TSTs across various cancers. The heatmap shows the top 30 TSTs with the highest expression frequency, and blue represents the number of tumor samples. g, Cancer specificity of TSTs. The median expression values of TSTs were calculated for each cancer type, and the heatmap color represents the normalized value of splicing junctions across different cancers. Abbreviation: NT, Tumor-adjacent normal tissue. TCGA cancer type abbreviations are found at https://gdc.cancer.gov/resources-tcga-users/tcga-code-tables/tcga-study-abbreviations.

Most TSTs are recurrent, with approximately 4882 TSTs appearing in at least 100 (>1%) samples (Fig. 3f). For instance, a TST derived from *XCL1* was expressed across several types of cancer, including bladder urothelial carcinoma (BLCA), esophageal carcinoma (ESCA), head and neck squamous cell carcinoma (HNSC), lung squamous cell carcinoma (LUSC), and other types of cancer (Extended Data Fig. 2a). Moreover, TSTs exhibit cancer-specific expression patterns (Fig. 3g), as exemplified by a TST from *ID4* specifically expressed in OV (Extended Data Fig. 2b). These cancer-specific features suggest the potential utility of TSTs as diagnostic biomarkers and therapeutic targets in cancer.

### TSTs are associated with poor prognosis and act as potential cancer drivers

We next explored the clinical significance of cancer patients with the presence of TSTs. A total of 3,130 TSTs were found to be associated with survival in at least one cancer (Extended Data Fig. 3a), and some of these TSTs were clinically relevant in multiple cancers (Extended Data Fig. 3b). To assess the degree of tumor-induced TST production, we developed a scoring system for each tumor sample on the basis of expression and frequency of TSTs (Fig. 4a, see Methods). Pan-cancer analysis of 9,555 tumor patients showed that higher TST level was significantly associated with worse overall survival (OS) and disease-specific survival (DSS), shorter progression-free interval (PFI) and disease-free interval (DFI) (Fig. 4b and Extended Data Fig. 3c). We found that TST level was associated with multiple clinical end points in several cancer types including pancreatic adenocarcinoma (PAAD), liver hepatocellular carcinoma (LIHC), brain lower grade glioma (LGG) and uveal melanoma (UVM) (Extended Data Fig. 3d).

**Figure 4.**
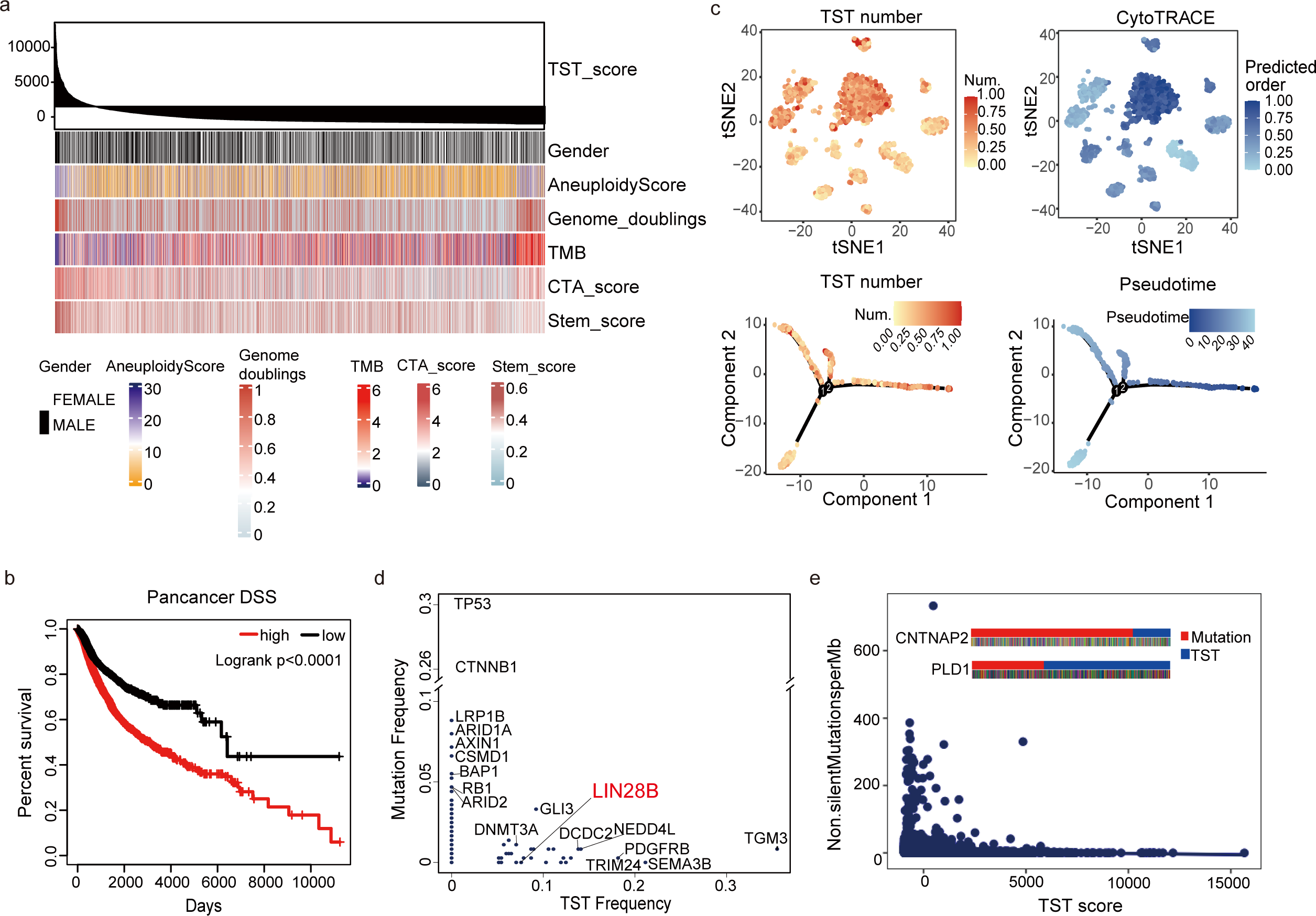
TSTs are associated with poor prognosis and act as potential cancer drivers. a, the relationship between transcriptional aberrancy level (TST score) and genomic and biological features. Each row represents a feature, each column represents a cancer sample, and they are sorted by TST score from high to low. The CTA_score row indicates the expression level of cancer/testis antigens. b, At pan-cancer level, TST score was associated with OS. c, The positive correlation between TST level and stemness validated with single-cell datasets. The upper panel shows tSNE plots depicting the distribution of TST number (0-1 normalized) and CytoTRACE score of single-cell samples. The lower panel shows pseudotime trajectory plots constructed by Monocle algorithm, showing that TST number distribution is consistent with cell differentiation trajectory. Color intensity indicates high or low number or score. d, The frequency of TSTs and mutations of cancer-related genes in liver cancer show opposite trends. The TST from LIN28B has been validated by our previous experiments and is marked in red. e, The TMB and TST scores also show opposite trends in pancancer; mutations and TSTs in CNTNAP2 and PLD1 are mutually exclusive (shown in the upper left corner of the figure).

We investigated the associations between TST score and other molecular features. The results showed that TST levels were positively correlated with genome aneuploidy and negatively correlated with tumor mutational burden (TMB) (Fig. 4a). Interestingly, a positive correlation was observed between TST level and stemness (Fig. 4a). The correlation between TST and stemness was calculated in 33 cancer types from TCGA. In 16 cancer types, TST levels showed a strong and significant positive correlation with stemness, with correlation coefficients ranging from 0.318 to 0.683 (Extended Data Fig. 4). To substantiate the correlation between TST level and stemness, we scrutinized a single-cell lung cancer dataset. 1969 cancer cells with a junction number greater than 15000 were retained for analysis using CytoTRACE, an algorithm that infers stem cell properties by scoring each cell based on its transcriptional diversity. As shown in Fig. 4c, Cells with more numerous TSTs displayed lower general differentiation (higher stemness, higher CytoTRACE score) compared to other cells (Fig. 4c). To reconstruct the acquisition of cell fate over time, we analyzed Monocle 2 to scRNA-Seq data, which provides pseudo-temporal ordering of cells relative to developmental progression (‘pseudotime’). It was also found that TST number changed along with cell differentiation trajectory (Fig. 4c). These results indicate that abnormal transcription of tumors is positively correlated with stemness.

To determine whether TSTs have functional effects, TST-derived genes were examined. We observed that some cancer driver genes such as *EGFR* and *ERBB2* can produce TSTs. It was further investigated whether TSTs can function as potential cancer driver events by focusing on genes affected by somatic mutations and/or TSTs. A clear pattern was observed wherein significantly mutated genes (SMGs) such as *TP53*, *CTNNB1*, *ARID1A*, and *RB1*, and significantly TST-produced genes (STGs) such as *TGM3*, *TRIM24*, and *LIN28B* were bifurcated into two groups in LIHC cohort (Fig. 4d), suggesting SMGs and STGs are mutually exclusive. Among STGs, LIN28B-TST was previously reported to encode a protein isoform with additional N-terminal amino acids and was critical for cancer cell proliferation and tumorigenesis^14^. This supported the notion that STGs could represent an independent catalog of genes with functionally important tumor-specific transcripts alterations in cancer. To explore whether lack of mutations in STGs is a general feature, we investigate the relationship between TST scores and TMB in the pan-cancer cohort. As expected, the results showed that samples with high TST levels had low TMB, while the low TST group had a higher level of mutation (Fig. 4e). When examining genes affected by somatic mutations and/or TST splicing such as *CNTNAP2* (one of the largest genes in the human genome, occupying 1.5% of chromosome 7) and *PLD1* (tumor-associated gene) in pan-cancer cohort, we confirmed this pattern of mutual exclusivity (Fig. 4e). The mutual exclusivity between TST and mutation suggests that TSTs may act as drivers similar to mutations in cancer development.

### TSTs are enriched in transposable elements derived transcripts with oncogenic function

Transposable elements (TEs) are major components of the human genome, comprising almost 50% of the total genomic DNA. We identified junctions derived from TE sites in both tumor and normal tissues. In total 154505 splicing TE junctions(spTEs) were detected in tumor tissues and 40737 were in normal tissues. The number of spTEs in cancer was much greater than in normal tissues (Fig. 5a), indicating that TE expression is activated in cancer. Strikingly, the TSTs were significantly enriched in TEs (tumor-specific TEs, tsTEs), particularly in the LTR class (Fig. 5b and Fig. 5c). Further analysis showed that tsTEs evolved much more recently than the background (Fig. 5d), possibly because young TEs are less affected by silencing mechanisms in tumors and have stronger transcriptional potential.

**Figure 5.**
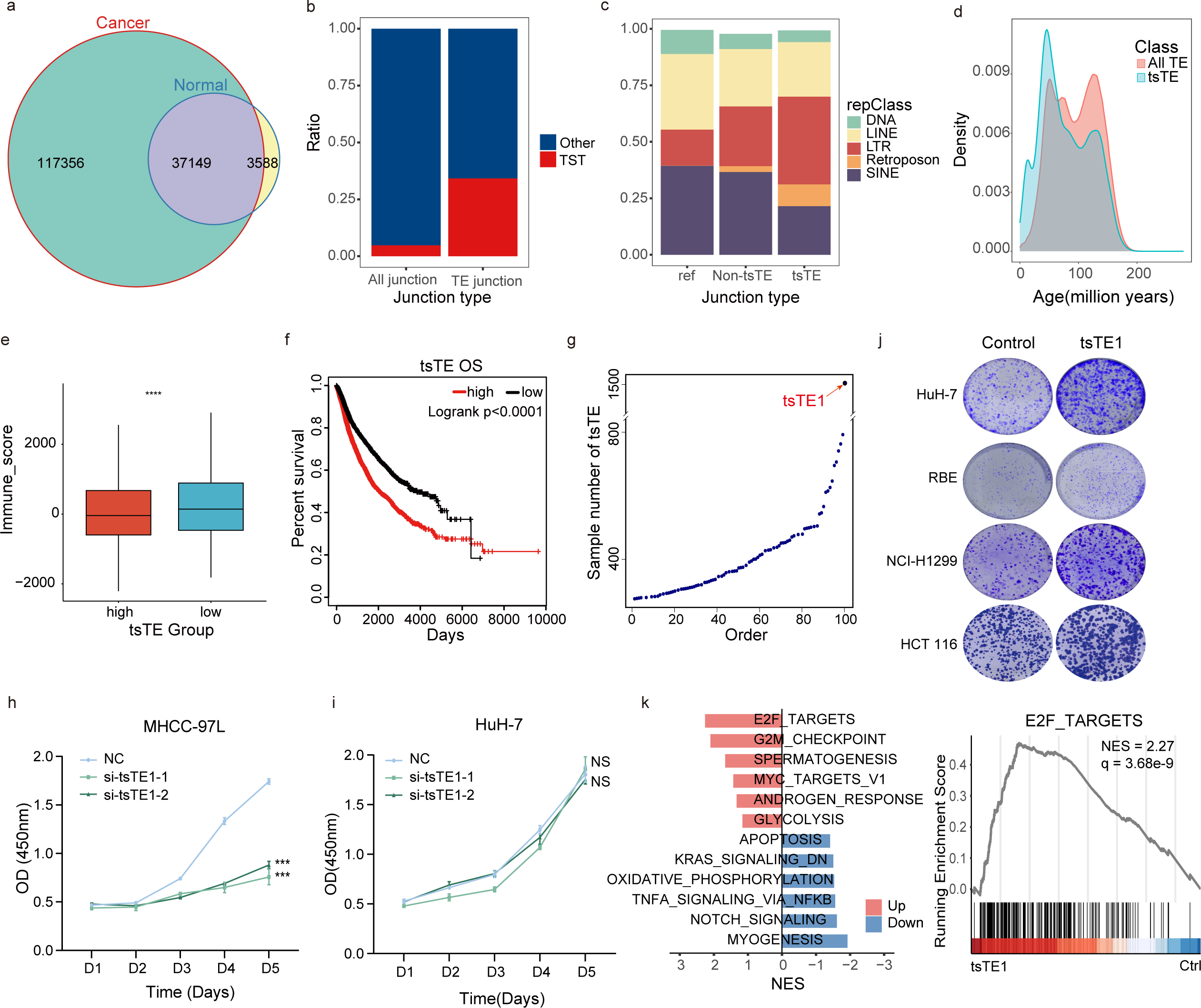
TSTs are enriched in transposable elements derived transcripts with oncogenic function. a, Splicing sites derived from TE identified in cancer samples and normal samples. b, TSJs are enriched in TE elements compared to all splicing junctions as background. c, The proportion of LTR elements and retrotransposons is increased in tsTE compared to all TE and non-tsTE, and LTR elements have the largest proportion among other TE categories. d, tsTE are generated from younger TE elements. The x-axis is the divergence age of TE (the smaller the value, the younger the TE), and the y-axis is the density of TE with corresponding divergence age. e, Boxplot showing immune score in samples with high levels of tsTE expression and low levels of tsTE expression. High, above median; Low, below median. p value from Wilcoxon test. f, Higher tsTE expression was correlated to worse overall survival in pancancer tumor samples. g, A large amount of tsTE identified in tumor samples are frequently expressed with tsTE1 showing the highest recurrence. h-i, Cell viability was inhibited following siRNA-mediated knockdown of tsTE1 in cells with high tsTE1 abundance (h) while barely affected in cells not expressing tsTE1 (i). j, Colony formation ability of various cancer cell lines were promoted by the stable overexprression of tsTE1. k, GSEA comparing transcriptome of HuH-7 cells overexpressing tsTE1 and control with the top 6 upregulated and downregulated pathways respectively shown (Left panel). Details of the enrichment were also presented for the top pathway associated with E2F targets (Right panel).

TEs are generally believed to activate the immune system by their transcript-derived components^15^. However, we found that high expression of tsTE was associated with lower immune activity in tumor samples (Fig. 5e) and correlated with poor prognosis (Fig. 5f), indicating a potential role for tsTE in promoting tumor development. Further analysis of tsTEs in cancer tissues revealed that a substantial number of tsTEs are frequently expressed (Fig. 5g). Among these, tsTE1 had the highest expression frequency in various cancer tissues (Fig. 5g and extended Data Fig. 5a). tsTE1 was a novel transcript located in cytoplasm (Extended Data Fig. 5b). RACE analysis confirmed that tsTE1 included a 511nt LTR sequence and was predicted to be a non-coding RNA (Extended Data Fig. 5c). Inhibition of tsTE1 expression by siRNA significantly inhibited the proliferation of cancer cells expressing tsTE1, while having no effect on cells that did not express tsTE1 (Fig.5h and Fig. 5i). Conversely, overexpression of tsTE1 using a lentivirus significantly promoted colony formation in various cancer cells (Fig. 5j and Extended Data Fig. 5d). Next, pathway analysis was performed using differentially expressed genes from samples with and without tsTE1 expression. The findings revealed that tsTE1 was significantly correlated with cell cycle pathways and may promote tumorigenesis by regulating cell cycle-related genes (Fig. 5k). These results suggest that tsTEs could have oncogenic functions in tumor development.

### TSTs represent a source of tumor neoantigens

Tumor neoantigens are currently important targets for cancer immunotherapy, but the discovery of neoantigens from tumor mutated proteins is limited. We investigated whether TSJs/TSTs derived tumor-specific peptides/proteins could potentially produce specific neoepitopes. We developed a workflow for predicting tumor neoantigens from RNA-seq (Extended Data Fig. 6). Among a total of 29,051 TSTs across 33 TCGA cancer types, 12,348 TSTs had the potential to encode proteins, accounting for 42.5% of all TSTs. We quantitatively assess the immunogenic potential of peptides across splicing junctions with coding capacity with a previously published score tool^16^. The immunogenic-related scores of 23,394 predicted translated-TSJ-derived peptides were ranked (mean: 0.263, range: 0-0.50) (Extended Data Fig. 7a). About half of peptides have strong affinity with MHC-I molecules in more than 5 patients (Extended Data Fig. 7b). Notably, TSJ introduced more potential neoantigens in tumors with low nsSNV-derived neoantigen load such as testicular germ cell tumors (TGCT), while mutation introduced more potential neoantigens in tumors with low TSJ-derived neoantigen load such as lung adenocarcinoma (LUAD) (Fig. 6a). In a further correlation analysis, we found a significant positive correlation between coding TSJ number (Extended Data Fig. 7c) and TSJ-derived antigen load, but a weak negative correlation between TMB and TSJ-derived antigen load (Extended Data Fig. 7d). Therefore, for cancer patients with a lack of non-synonymous mutation-derived neoantigens, TSJ-derived antigens may offer potential therapeutic targets.

**Figure 6.**
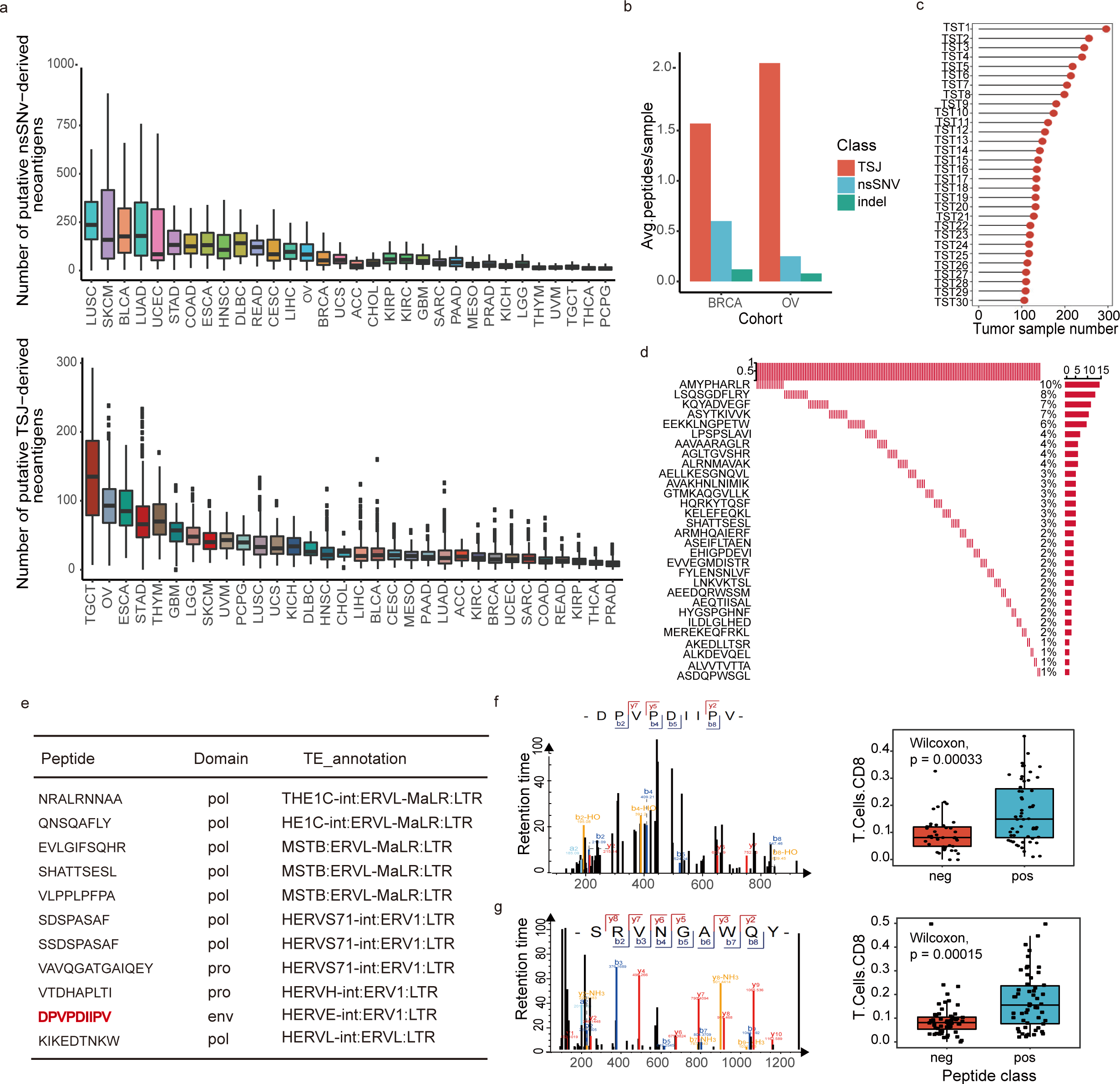
TSTs represent a source of tumor neoantigens. a, Neoantigen load of mutation origin and TSJ origin in 32 TCGA tumors. LAML (lymphoma) is a malignant blood tumor and is not included. b, Number of neoantigens from TSJs, somatic SNVs and indels confirmed by CPTAC in BRCA and OV. c, TSTs generating immunopeptides can be present in multiple samples. The top 30 most frequent TSTs are shown here. d, TSTs-derived neoantigens can be detected repeatedly in multiple immunopeptidomics samples. Each row in the waterfall plot represents a neoantigen identified by immunopeptidomics and each column represents an immunopeptidomics sample. The frequency of each peptide is shown on the right. e, Immunopeptide sequences and family information derived from ERV elements. f, Schematic diagram of immunopeptides from ERV-derived neoantigens and immunopeptidomics spectra (left), and comparison of CD8+ T cell infiltration level between samples with or without tsTE peptides (right). g, Schematic diagram of immunopeptides from non-TE regions and immunopeptidomics spectra (left), and comparison of CD8+ T cell infiltration level between samples with or without this peptide (right).

To validate the expression of these predicted peptides, we used MS-GF+^17^ to search mass spectrometry (MS) data in ovarian cancer (OV) and breast invasive carcinoma (BRCA) samples available from the Clinical Proteomic Tumor Analysis Consortium (CPTAC) project, and identified a total of 37 neoepitopes derived from TSJs (Supplementary Table 4). Compared to non-synonymous single nucleotide variants (nsSNVs) and insertions/deletions (indels), TSJs introduced a higher number of CPTAC-confirmed neoantigens per sample in both OV and BRCA (Fig. 6b). Although CPTAC proteomic data confirmed the expression of peptides derived from TSJs, they were not able to demonstrate whether these peptides were processed and presented on major histocompatibility complex (MHC). To validate this, we conducted an analysis on 352 immunopeptidome samples of different cancer cells or tissues (Supplementary Table 4). Due to limited detection sensitivity of immunopeptidome and TSJs may cause in-frame deletions of functional protein domains or generate novel reading frames through frameshifts, we identified neoepitopes derived from TSTs. Altogether, 257 TST neoantigens were confirmed to be presented by MHC-I molecules, and both TSTs and TST-derived immunopeptides could be reproducibly detected in corresponding samples (Fig. 6c and Fig. 6d). The TST-derived peptides and UniProt annotated peptides had similar peptide length distributions, with the 9-mer peptides being most frequently presented by MHC-I molecules (Extended Data Fig. 7e), and the TST-derived 9-mer peptides had similar binding motifs with UniProt annotated peptides (Extended Data Fig. 7f).

Among these TST-derived immunopeptides, some peptides from TEs are derived from validated endogenous viral elements (EVEs) in the gEVE database^18^. These EVEs were identified by processing both RepeatMasker annotations and conserved known motifs from viral proteins such as Gag and Pol. ERVL-MaLR and ERV1 elements show selectivity in providing immunopeptides presented by MHC-I molecules (Fig. 6e). For example, DPVPDIIPV is a novel epitope produced by RNGTT-derived TST containing ERV1 elements, which was confirmed by mass spectrometry-based immunopeptidome analysis and induced increased levels of CD8+ T cell immune infiltration (Fig. 6f). Another example is SRVNGAWQY produced by TST from non-TE regions, which also causes differences in CD8+ T cell infiltration levels (Fig. 6g). These results suggest that TST-derived peptides are credible and may serve as a valuable source for neoantigen discovery.

We next sought to examine whether TST-derived neoantigen load was associated with clinical response to immune checkpoint blockade (ICB). To test this, we analyzed data from one cohort of metastatic gastric cancer patients receiving pembrolizumab treatment. We found patients with clinical benefit had a higher number of TST neoantigen load than patients without benefit (Extended Data Fig. 8a). The expression level of immune checkpoint molecules was considered one of the indications for immune checkpoint inhibitor therapy. We compared the expression of 16 literature-curated immune checkpoint molecules between high and low neoantigen load patients. The expression of *PD-1*, *CD276*, *HAVCR2*, *LAG3*, *SIGLEC7* and *SIGLEC9* was significantly higher in high neoantigen load group (Extended Data Fig. 8b). In addition, we found that six patients had a higher level of TST-derived neoantigen load, four of whom achieved complete or partial remission (CR/PR) after receiving immunotherapy (Extended Data Fig. 8c). Conversely, among 17 patients with moderate or low TST-predicted neoantigen loads, immunotherapy was not effective in 12 cases (Extended Data Fig. 8c).

### RNA splicing junctions in extracellular vesicles for cancer diagnosis

Extracellular vesicles (EVs) are small heterogeneous vesicles that are secreted by virtually all cells and are abundant in all body fluids. They carry various molecular cargo, including RNA, and act as a reliable liquid biopsy for disease diagnosis^19^. We performed EV RNA-Seq analysis of 1128 samples from four different body fluids. We identified a total of 472,193 linear-splicing sites and 121,816 back-splicing sites in EVs. We found 46.83% (349,613/746,461) of the tissue/cell expressed linear-splicing junctions are detectable in EVs (Fig. 7a). RNA splicing junctions in EVs and tissues have a similar genomic distribution (Fig. 7b), and circRNAs are significantly enriched in EVs (Fig. 7c).

**Figure 7.**
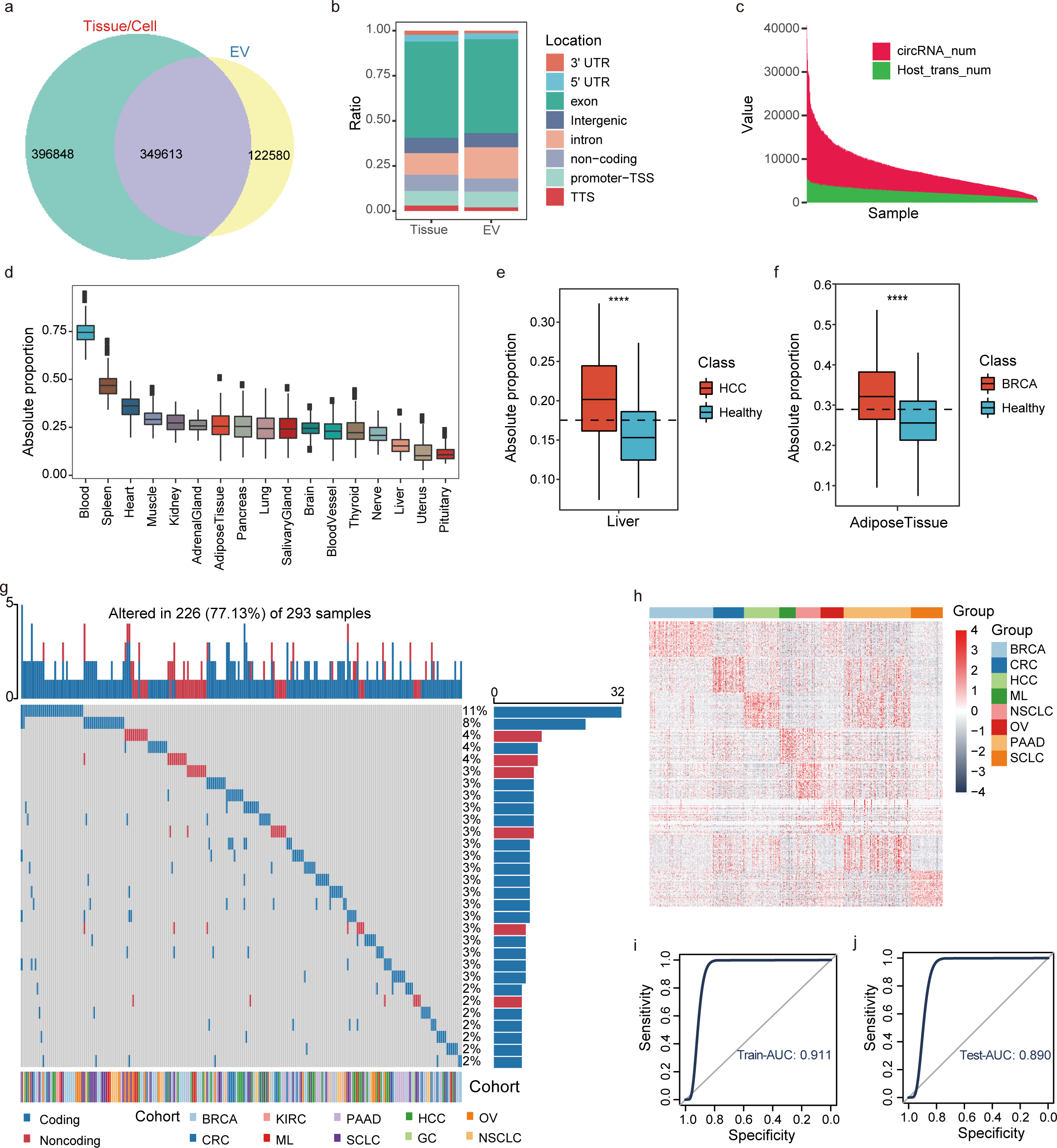
RNA splicing junctions in extracellular vesicles for cancer diagnosis. **a,** Splicing junctions detected separately and commonly in tissue/cell and EVs. **b,** Genomic location distribution of splicing sites in tissue and EVs. **c,** The distribution of back-splicing (circRNA) and linear junction numbers in EVs. **d,** Proportion of normal tissues. **e,** Difference of EVs derived from liver tissue between HCC and healthy samples. **f,** Difference of EVs derived from adipose tissue between BRCA and healthy individuals. **** represents p-value < 0.0001 by Wilcoxon test. **g,** TSJ expression in plasma EVs. The waterfall plot shows the top 30 splicing sites with highest expression frequency. The top shows the number of TSJs expressed in each sample, the right bar plot shows the expression frequency of each TSJ, the bottom shows the TSJ category and tumor type. **h,** Cancer-specific splicing sites. The performance of the SVM model to distinguish different types of cancer in training set(**i**) and test set(**j**) based on cancer-specific splicing sites.

It has been demonstrated that specific cargo of EVs is reflective of donor cell origin and its physiological state^20^. We performed a tracking analysis with RNA splicing sites for investigating the origins of plasma EVs at tissue level. Fig. 7D showed the deconvoluted tissue-proportion analysis of EV sources based on the expression of tissue-specific splicing sites (TSS) in the 1,076 individual plasma samples. We found that plasma EVs were mainly contributed by the spleen tissue in addition to blood (Fig. 7d). When considering cancer types, we observed that TSS of the liver were enriched in liver cancer compared to control samples (Fig. 7e), while TSS of adipose tissue were enriched in breast cancer compared to control samples (Fig. 7f). Furthermore, we observed that tumor-exclusive splicing junctions-parent genes were enriched for metabolic and tumor-related pathways, while immune-related signatures were exclusively enriched in EVs (e.g., T cell and B cell receptor signaling pathways) (Extended Data Fig. 9a).

To investigate whether RNAs with specific splicing sites can be used as cancer biomarkers, we first examined the expression of TSTs in plasma EVs. Interestingly, TSTs could be detected in EVs and were expressed in multiple samples (Fig. 7g). Nevertheless, the overall frequencies of TSTs were low (293 out of 747 cancer samples expressed TSTs) (Fig. 7g), possibly due to insufficient sequencing sensitivity. Additionally, we identified 194 specific RNA junctions that had significantly different levels between patients and healthy individuals (Extended Data Fig. 9b), of which 178 (17 were from tumor-associated genes) were upregulated in cancer and only 16 downregulated (Extended Data Fig. 9b and Extended Data Fig. 9c). For different types of cancer, up-regulated splicing sites were varied. For example, the splicing sites from APOH and HRG genes were specifically expressed in liver cancer (Extended Data Fig. 9d), while ETV4 and EAF2 are specifically expressed in pancreatic cancer (Extended Data Fig. 9e) and breast cancer EVs (Extended Data Fig. 9f), respectively. We therefore identified cancer-specific splicing sites (Fig. 7h) and investigated whether they can distinguish cancers. The EV datasets, encompassing seven cancers, was partitioned into training (70%) and validation (30%) subsets. A multi-class Support Vector Machine (SVM) model was employed to construct a splicing classifier using these cancer-specific junctions. The efficacy of the splicing classifiers was assessed via Receiver Operating Characteristic (ROC) analysis, focusing on the Area Under the Curve (AUC). The diagnostic ROC curves demonstrated outstanding AUC values: 91.1% for the training set (Fig. 7i) and 89.0% for the validation set (Fig. 7j), underscoring the robust predictive power of the model. These results suggest that RNA splicing sites in EVs are rich sources of diagnostic targets for different types of cancer.

## Discussion

In this study, we collected 34,775 RNA-seq samples from diverse human tissues and cell types, and obtained a comprehensive and robust repertoire of RNA splicing landscape in the human transcriptome. It is noteworthy that nearly half of these splicing junctions had remained un-annotated and largely overlooked by previous studies. We further discovered 29,051 TSTs from 32,848 tumor-specific splicing sites. Our investigation revealed a significant positive correlation between TST levels and stemness, while showing mutual exclusivity with mutations. Moreover, we explored generating novel cancer antigens through TSTs, finding that TSTs are translated into tumor-specific proteins containing peptides with MHC presentation potential. At the splice site level, we demonstrated the utility of RNA present in plasma extracellular vesicles for identifying cancer biomarkers and tracking tissue/cell sources. This study is the first to reveal RNA splicing junction landscape and discover a large number of TSTs with potential functions in pan-cancer.

We investigated the RNA-seq data at splice junction level by ASJA, a tool that identifies splicing sites from reference-based assembly transcripts, resulting in more reliable data. Most of the current computational tools for alternative splicing analysis are based on the reference genome and allows for more accurate isoform detection^21,22^, which may result in loss of large amounts of putative intergenic transcripts. Compared to reference-based assembly, de novo assembly approaches have the advantage of detecting genes in the absence of a reference genome while being associated with difficulties in correction of read errors, calling for the merging of all assembly transcripts before quantification and comparison^22^. Given that expression of assembly transcripts is often inconsistent across different samples, it is challenging to handle different types and large proportions of sequencing data. ASJA is an effective approach by focusing on the junctions of reference-based assembly transcripts (31462970). Compared to junction analysis, which is performed using mapped reads, ASJA extracted junctions from the assembly transcripts, resulting in junction data that are more reliable at the source. Analysis of such junctions in one sample produces a unique junction position and expression level, allowing for direct comparisons to be performed across different samples. Moreover, all the junctions are included. Therefore, almost half of the identified splicing sites are unannotated in this study. Chromatin open signals near the transcription start site of unannotated transcripts indicate the authenticity of their source transcripts. In addition, we observed many non-canonical splicing events that occurred within genomic regions including intergenic regions, exons, introns and untranslated regions (UTRs) in our study. Some non-canonical splicing sites have been recently studied. For example, Zhang et al. found that 3’ UTR splicing is commonly present in cancer and is associated with poor prognosis^10^. Wang et al. confirmed that exitron splicing can affect the function of cancer driver genes and promote tumor progression^11^. These non-canonical splicing events escape cellular quality control mechanisms, and their role as a source for new molecular functions during evolution will remain a fascinating subject of research.

Tumor-specific transcripts (TSTs) containing tumor-specific splicing events are promising sources of novel therapeutic targets in cancer. By comparing splicing junctions between cancer and normal samples, we discovered 29,051 TSTs in pan-cancer, nearly half of which were derived from intergenic regions. The extent of aberrant splicing was positively associated with stemness, indicating TSTs may participated in establishing the stem-like properties that typify the more aggressive nature of neoplastic cells. Interestingly, we found that TST and TMB were mutually exclusive in cancer especially in driver genes, which is significant as mutation analysis is an established approach for identifying driver genes in cancer. Similar to *LIN28B* which has been validated to enhance cancer cell proliferation and tumorigenesis^14^, we have also identified a tumor-specific transcript produced by *MARCO*, namely MARCO-TST^13^. This transcript plays a crucial role as an oncogenic transcription factor and therapeutic target, inducing metabolic dysregulation and hypoxic microenvironment in triple-negative breast cancer^13^. Therefore, the analysis of TSTs has the potential to reveal previously undetected cancer driver genes, highlighting the need for further development of both computational and experimental technologies for detecting and characterizing these driver splicing events. Furthermore, we discovered that many splicing events are introduced by TEs, which constitute up to half of the human genome. We found TSTs to be enriched in TEs and experimentally validated cancer-promoting function of tsTE1. This indicates that cancer utilizes degenerated TEs to generate new exons, producing TSTs that promote cancer development and progression.

Immunotherapy has revolutionized cancer treatment and brought renewed hope to patients with hard-to-treat and invasive cancers by significantly improving survival rates for certain types of cancer^23^. To achieve safe and effective immunotherapy, it is crucial to discover highly cancer-specific and immunogenic tumor antigens (TA)^24^. Currently, TAs are primarily derived from somatic mutations in protein-coding regions that generate neoepitopes^25^. However, identifying effective targets for cancers with low mutational burdens can be challenging as only a few somatic mutations can change the protein sequence and confer immunogenicity. The TSTs with protein coding ability may provide valuable insights for identifying novel TA and developing effective immunotherapies. We analyzed the contribution of TSTs to potential neoepitopes to substantiate the hypothesis regarding their significance in the immune response to cancer. An analysis pipeline was developed to go from HLA typing to the predicted translations of peptides from TSTs, and confirmation using MS data and immunopeptidome data. We found that TSJs introduced more potential neoepitopes in tumors with low nsSNV-derived neoantigen load. TSJs also introduced a higher number of CPTAC-confirmed neoepitopes per sample than nsSNVs and indels in both OV and BRCA. Using 352 immune peptidome data, we identified 257 peptides derived from TSTs. Previous studies have indicated that the majority of MHC-binding peptides are derived from non-coding genomic regions, which mainly comprise transposable elements such as ERVs known to generate tumor-associated antigens and promote immune response^26-28^. Among the extracted immune peptide segments, we identified 11 peptides originating from tsTE sources containing ERV elements, confirming that both non-TE and ERV-derived peptides can elicit differential expression of CD8+ T cells. Although this study used MS-based proteomics analysis to validated the expression of peptides and MS-based immune peptide profiling data to identify peptides presented by MHC, it is still limited by the technology’s sensitivity, and the use of more sensitive proteomic techniques will facilitate the identification of novel neoepitopes. Additional immunological experiments are needed to advance the development and clinical translation of the TST-based vaccines.

Our investigation has revealed the presence of abundant splicing events within circulating EVs. EVs contain various biotypes of RNA, and previous studies have demonstrated their potential as promising biomarkers for disease diagnosis^19,29^. Notably, our study demonstrates that nearly half of the RNA splicing junctions found in tissue or cells could also be detected in EVs, with a subset showing tissue-specificity and thus serving as potential EV biomarkers for origin tracing. Importantly, we also detected TSJs in the plasma EVs of cancer patients, highlighting the potential of EV-encapsulated RNA splicing as a prognostic marker for cancer. However, due to the stringent criteria for identifying TSJs in tissues and the limited sequencing depth of exLR-seq, only a small fraction TSJs were recovered from plasma EVs, resulting in insufficient coverage of the samples. Despite these limitations, our analysis of differential targets and model construction revealed that cancer-specific splicing junctions could distinguish various types of cancer with AUC values exceeding 86%. This finding underscores the potential of EV RNA splicing analysis as a practical, noninvasive approach for molecular-level diagnosis and prognosis using human plasma EVs.

In summary, our study comprehensively analyzed RNA splicing sites in the human transcriptome and identified numerous TSTs with potential functions in cancer. The splicing landscape we characterized provides valuable insights into the lineage-specific and expression patterns of splicing events in cancer. The identification of TSTs has significant implications for cancer biomarker discovery, identification of cancer driver genes, and detection of neoantigens, which opens up new avenues for cancer diagnosis and therapy.

## Materials and methods

### Sample acquisition

We retrieved raw RNA-seq data from TCGA and CCLE databases (http://cancergenome.nih.gov and https://portals.broadinstitute.org/ccle, respectively). Linear junction data in normal tissues were obtained from the Genotype-Tissue Expression (GTEx) portal(https://gtexportal.org). The raw data of single cells were downloaded from PRJNA591860^30^, which included 4,936 samples from six biopsy sites (adrenal, brain, liver, lymph node, lung and pleura). We performed EV RNA-Seq of 1128 samples from four different body fluids by using exLR-seq as we previously reported^31^. The detailed samples were listed in Supplementary Table. 1. Protein mass spectrometry data from the CPTAC cohort are available at the CPTAC data portal: https://cptac-data-portal.georgetown.edu/cptacPublic/^32^. Multiple types of immunopeptidomics datasets from independent sources were used in this study and can be found at PRIDE (http://www.ebi.ac.uk/pride/archive/) under IDs: PXD007635, PXD013649, PXD014017, PXD027182, PXD009925, PXD022150, PXD06939, PXD022950, PXD013831.

### Identification and quantification of RNA splicing junctions

The linear splicing junctions from TCGA, CCLE cancer cell line, EVs and single-cell samples were extracted and quantified using ASJA^33^. Briefly, RNA-seq reads were aligned to the hg38 reference genome by using the STAR two-pass alignment algorithm^34^, followed by transcriptome assembly using StringTie with human GENCODE V29 annotation file as reference. ASJA extracts linear splicing junctions based on the assembly results and obtains splice sites as well as standardized expression values. We obtained linear junction data from datasets stored at the Genotype-Tissue Expression (GTEx) portal, and the raw count were normalized using the formula: raw_counts/total_counts × 10 000 000 (total_count is the sum of all splicing counts in the sample), with the scale factor being the same as that adopted by ASJA. Junctions that were present in fewer than five samples of each data cohort were removed.

### RNA splicing junction annotation

The gene boundaries were sourced from the GENCODE database. The coordinates of each splicing junction were then annotated with the nearest gene name utilizing bedtools closest and bedtools intersect^35^. We obtained a pre-annotated splicing set by extracting splicing events from the GENCODE V29 reference GTF file. Any excluded junctions were marked as unannotated. To classify each splicing event, we first extracted and dismissed constitutive splicing sites. Next, we employed a tailored script to identify five types of alternative splicing junctions based on donor and acceptor coordinates. However, single splicing sites do not cater to exon skipping or retained introns measurements. Consequently, they were not included there. The remaining splicing sites were categorized into three categories according to their genomic locations: intergenic, intronic, exonic (cds and UTR).

### Validation of splicing junctions identified from RNA-seq

Glinos et al. used Oxford Nanopore Technologies’ cDNA-PCR protocol to generate long-read RNA-seq data from 88 GTEx tissues and obtained transcripts with FLAIR^36^. We downloaded the transcript GTF file from the GTEx website (https://gtexportal.org/home/datasets) and extracted linear splice sites using a Python script. Additionally, direct RNA sequencing data for Huh7 and HCT116 cell lines were performed. We aligned direct RNA sequencing data for Huh7 and HCT116 cell lines to the hg38 reference genome using minimap2 within FLAIR and identified linear splice sites with 2passtools^37^. The RNA-seq data for Huh7 and HCT116 were processed using the ASJA pipeline. Finally, we calculated the proportion of RNA-seq splicing junctions identified from RNA-seq which also can be identifed from long-read RNA-seq data.

To assess the authenticity of transcripts containing unannotated splice sites, we downloaded the ATAC-seq bigWig files from TCGA (https://gdc.cancer.gov/about-data/publications/ATACseq-AWG/) to evaluate chromatin accessibility near transcript regions. From all TCGA cancer samples, we selected the most highly expressed transcript with unannotated splicing junctions and identified its transcription start site (TSS) location. The chromatin accessibility around this region was assessed by evaluating the enrichment of ATAC signals within 5kb of the TSS using ‘computeMatrix’ and ‘plotHeatmap’ commands from deptools^38^.

### RNA splicing efficiency evaluation

The splicing frequency and efficiency of both protein-coding genes and lncRNA genes were evaluated. Splicing frequency represents the number of splicing events generated by these gene types. Given the correlation between splicing frequency and exon count, we further analyzed splicing efficiency by dividing the gene’s respective splicing number by its exon number.

### Identification of tissue-specific genes and splicing junctions

We leveraged splicing and gene expression data obtained from GTEx to identify tissue-specific splicing events and genes. GTEx encompasses a total of 17,382 samples across 30 tissues, albeit with unbalanced sample sizes per tissue ranging from 21 in the bladder to 3,326 in the brain. Our study utilized SPECS^39^, which offers a new nonparametric specificity score optimized for biological replicate expression matrices of varying sample sizes. By implementing SPECS, we calculated tissue-specificity scores for each splicing junction and gene. In this regard, we defined splicing events with SPECS scores exceeding 0.95 as tissue-specific splicing, while genes with SPECS scores exceeding 0.95 were deemed as tissue-specific genes.

### Identification and characterization of tumor-specific transcripts

To identify tumor-specific splicing junctions (TSJs), splicing expression were compared between tumor (TCGA) and normal (TCGA plus GTEX) samples across various cancers. Splicing junctions that met the following criteria were identified as TSJs: NT_max = 0 & nTE_max = 0 & Freq_cancer > 5%; Tumor_median > 10*NT_median & Tumor_median > 10*nTE_max & Freq_cancer > 5%, where Tumor_median represents the median expression value in tumor samples, NT_max is the maximum expression value in TCGA adjacent normal samples, nTE_max is the maximum expression value in GTEx samples (excluding testis tissue), and Freq_cancer reflects the frequency of expression in cancer for the splicing junction. Transcripts containing TSJ sites were classified as Tumor-specific transcripts (TSTs). In order to simplify and standardize our analysis, we selected TST from TCGA with the highest expression among 9,575 tumor samples as the tumor-specific transcript corresponding to the TSJ. The expression value of TSTs was defined as the maximum expression value among the TSTs including TSJs. We obtained the donor and acceptor positions of TSTs and employed the annotationPeaks.pl command of the HOMER program^40^ to annotate them to promoter regions (within 2kb of known TSS), intergenic, intronic, exonic, 3’ UTR, and 5’ UTR regions.

### Correlation of TST level with clinical and molecular features

Clinical outcome endpoints data and patient gender and tumor stage data were obtained from the TCGA Pan-Cancer Clinical Data Resource^41^. Stemness scores were obtained from previously published datasets^42^. Additionally, we obtained information on tumor mutational burden (TMB) and non-ploidy score for all TCGA cancer samples from Taylor et al.’s publication^43^. In this study, we utilized TCGA data on gene expression and gene mutations from 33 types of tumors. Specifically, we downloaded the FPKM expression matrix for genes and the mutation matrix for “MC3” gene level from UCSC Xena (https://xenabrowser.net/datapages/).

To compare the level of TST transcription across different samples, TST score is calculated based on the expression value and frequency of identified tumor-specific transcripts (TSTs). The formula of TST score is as follows:

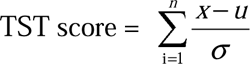

where i is the number of TSTs in the sample, x is the expression value of this TST in the sample, u is the mean expression value of TSTs across all TCGA cancer samples, and σ is the standard deviation of TST expression across all TCGA cancer samples. In other words, each TST is standardized to a z-score at the pan-cancer level, and the TST score is defined as the sum of these z-scores.

All TCGA cancer samples were sorted based on their TST scores and investigated whether any specific genomic/molecular/clinical features were associated with aberrant splicing in tumors. For summarizing the data, heatmaps was generated using the R Bioconductor package ComplexHeatmap^44^. To assess the correlation between TST scores and stemness, ggscatter function was applied in ggpubr to calculate Pearson correlation coefficients and associated p-values. To validate the positive correlation between TSTs and stemness, we utilized single-cell expression data derived from human lung tissue. CytoTRACE algorithm^45^ was applied to calculate the CytoTRACE scores for malignant cells. CytoTRACE scores range from 0 to 1, while higher scores indicate higher stemness (less differentiation) and vice versa. Pseudo-time analysis was performed with Monocle v.2.14.0 35^46^. To evaluate the impact of aberrant splicing on immunity in tumors, a list of immune checkpoint molecules was literature-curated and R package circlize^47^ was udsed to create circos plots. Lists of oncogenes and tumor suppressor genes were obtained from the ONGene^48^ and TSGene^49^ databases and referred to them as cancer-related genes.

### RNA splicing junctions in transposable elements

Transposable element annotations were from UCSC Genome Browser (RepeatMasker), and following information were obtained on each repeat: Class, Family, Subfamily, Divergence, coordinates. The bedtools (bedtools intersect) were used to extract the splicing junctions that overlap with TE regions. annotatePeaks.pl from Homer was performed to obtain genomic locations (intron, exon, 3′UTR, 5′UTR, intergenic, other) for each individual TE. Repeat age was calculated using percentage of divergence for human repeats: Divergence/(2.2 ∗ 10^-9^), following the formula from this article^50^.

### Cell culture

The cell lines used in this study, including HEK293T, MHCC-97L, HuH-7, RBE, NCI-H1299 and HCT 116 cells, were cultured in DMEM, RPMI1640 or McCoy’s 5A medium containing 10% fetal bovine serum, 100mg/ml penicillin, and 100U/ml streptomycin and incubated at 37□ with 5% CO_2_ in the humidified incubator.

### RNA preparation and quantitative RT-PCR

Total RNA was extracted with TRIzol reagent (Life Technologies) and subsequently transcribed into complementary DNA (cDNA) using Evo M-MLV RT Premix for qPCR kit (Accurate Biology). Quantitative realtime PCR (qRT-PCR) analyses were then performed using SYBR Green Premix Pro Taq HS qPCR Kit (Accurate Biology) on 7900HT Fast Real-Time PCR System (Applied Biosystems). Relative gene expression was calculated by 2^-ΔΔCt method normaized to the expression of ACTB. Each qRT-PCR assay was performed in triplicate and the primers used are listed in Supplementary Table 5.

### Rapid amplification of cDNA ends (RACE)

We applied 5’- and 3’-RACE assays to amplify the full length of tsTE1 transcript from the cRNA of MHCC-97L cells using the SMARTer™ RACE cDNA Amplification Kit (Clontech, Mountain View, CA, USA) following the manufacturer’s protocol. The gene-specific primers used for RACE analyses were listed in Supplementary Table 5. The PCR products were subsequently cloned to the pMD19-T vector (TaKaRa) for Sanger sequencing.

### Vector construction and transfection of cell lines

The full-length of tsTE1 was amplified from the cRNA of MHCC-97L cells as stated in Rapid amplification of cDNA ends (RACE) subsection, and then cloned into pWPXL vector (Addgene #12257) generating pwpxl-tsTE1 plasmid for overexpression assay in vitro.

siRNAs were synthesized by Ribobio (Guangzhou, China) and the sequences are provided in Supplementary Table 5. Cells were transfected at a final concentration of 20 nM using Lipofectamine RNAiMax (Life Technologies) according to the manufacturer’s instructions. After 48 hours cells were harvested for further analysis. For overexpression assay, the pwpxl-tsTE1 plasmid and packaging vectors (pMD2.G and psPAX2) were transiently co-transfected into HEK293T cells using Hieff Trans® Liposomal Transfection Reagent (Yeasen Biotech) to generate lentivirus particles collected after 48 hours. The target cells were subsequently treated with lentivirus particles and 8 µg/mL polybrene for 16-24 hours. The efficacy of siRNA interference and tsTE1 overexpression were both verified by qRT-PCR.

### Cell proliferation and colony formation assays

Cells were seeded into 96-well culture plates at the density of 3000 cells per well after transfected with siRNA oligos and cultured for 5 days with the viability measured daily using the Cell Counting Kit-8 (CCK8) (TargetMol) according to the manufacturer’s protocol. For the colony formation assay, cells transfected with lentiviruses expressing GFP or tsTE1 were plated into 6-well plates with 1000 cells per well. After 10-14 days, the colonies were fixed and stained with 1% crystal violet for 15 minutes. Cell colonies were imaged and counted by ImageJ software (1.54d) with default settings.

### Subcellular fractionation and RNA-seq analysis

Cytoplasmic and nuclear RNA were isolated with Cytoplasmic lysis buffer composed of 20mM Tris-HCl (pH=7.5), 10mM NaCl, 0.05% NP-40 and 2mM MgCl2. Briefly, MHCC-97L cells were scraped from a 6cm culture dish and next washed twice with cold PBS. Cell pellets collected were lysed in 300ul Cytoplasmic lysis buffer, pipetted and subsequently incubated on ice for 2 minutes with occasional gentle inversion. The cell suspension was then centrifuged at 4°C 3000rpm for 3 minutes. The resulted supernatant and pellet containing cytoplasmic and nuclear RNA respectively were separated into different tubes and used for RNA extraction.

Total RNA of HuH-7 cells transfected with lentiviruses expressing tsTE1 or GFP were extracted and treated with mRNA Capture Beads (Yeasen Biotech) to enrich polyA+ RNA before constructing the RNA-seq libraries. RNA-seq libraries were prepared using mRNA-seq Library Prep Kit for Illumina (Yeasen Biotech) following the manufacturer’s instructions. Sequencing reads from RNA-seq data were aligned using the spliced read aligner HISAT2^51^, which was supplied with the Ensembl human genome assembly (Genome Reference Consortium GRCh38) as the reference genome. Gene expression levels were calculated by the FPKM (fragments per kilobase of transcript per million mapped reads). Gene Set Enrichment Analysis^52^ (GSEA) was performed at the pre-ranked mode evaluating the enrichment of gene sets retrieved from the Molecular Signatures Database (MSigDB).

### HLA typing with seq2HLA

HLA class I four-digit types of 8,915 out of 9,599 TCGA tumor samples were obtained from Thorsson et al^53^. For the remaining 45 metastatic gastric cancer samples used in this study, seq2HLA version 2.2^54^ was used to infer the HLA class I four-digit types of patients from RNA-seq data.

### Prediction of tumor neoantigens derived from TSJ/TSTs

We developed a pipeline to predicte TSJ/TST-derived neoantigens (Extended Data Fig. 5). In brief, the pipeline is comprised of the following steps:

1. Sequencing data and pre-processing: Raw sequencing reads were quality-inspected using FastQC, and HLA typing was performed using the Seq2HLA tool on RNA-Seq data. Reads were further filtered and trimmed for quality using Trimmomatic 0.33^55^.
2. TSJ/TST identification: TSJs and TSTs were extracted as previously described. We retrieved all TST sequences in the genome using the getfasta function from bedtools (v2.30.0), with the TST transcript GTF file processed into BED format. CPAT software^56^ was utilized to evaluate the protein-coding potential of TSTs, and only those with a CPAT score greater than 0.364 were selected for further analysis. The ORF sequences of these selected TSTs were extracted and translated into amino acid sequences using the Biopython package^57^. TSJ-derived candidate peptides were defined as ten adjacent amino acids on both sides of the TSJs which parent TSTs with coding ability.
3. Neoantigen prediction: TST protein sequences were digested into small peptide fragments using a custom Python program. To simplify the analysis and remove the influence of peptide length, we selected 9-mer MHC-I-binding peptide segments for further analysis, as previous studies have reported that MHC-I preferentially binds to 9-mer peptide segments^58^, which has also been confirmed in immunopeptidome analysis. We prepared patient-specific peptide libraries by dividing full-length TST peptide segments into overlapping 9-mer peptide segments. Peptides derived from UniProt were removed, and the remaining peptides were used for further analysis. Similarly, 9-mer peptides derived from TSJs were obtained. NetMHCpan (v3.0)^16^ was utilized to predict the binding strength of TSJ/TST-derived peptides to patient-specific HLA alleles, and only those predicted to be strong binders were retained for subsequent analysis.

### Identification of TST-expressed peptides in CPTAC

Proteomic data from 32 samples of breast and ovarian cancer in The Cancer Genome Atlas (TCGA) were obtained from the Clinical Proteomic Tumor Analysis Consortium (CPTAC) data portal. For each tumor sample, a polypeptide database was constructed, consisting of mutant peptide sequences with ten adjacent amino acids on both sides of the TSJs concatenated with the UniProt human proteome. OpenMS software^59^ was utilized to identify the polypeptides within each sample’s database through the following steps:

1. decoy sequences were added to the database to manage false discovery rates using the command: DecoyDatabase -in <in.fasta> -out <db.fasta>;
2. MS-GF+ software was used to search the corresponding CPTAC mass spectrometry dataset against the polypeptide database with the following parameters: MSGFPlusAdapter -ini <config.ini> -in <spectra.mzML> -out <out.idXML> -database <DB.fasta> -executable MSGFPlus.jar -java_memory 20,000 -threads 16;
3. the mapping of peptides to proteins was refreshed, and target/decoy information was added using PeptideIndexer with the command: PeptideIndexer -in <out.id-XML> -fasta <DB.fasta> -out <pi_out.idXML> -allow_unmatched -enzyme:specificity ‘semi’;
4. peptide identification files from multiple runs were merged using IDMerger with the command: IDMerger -in <pi_out.idXML_files> -out <merged.idXML>;
5. we controlled for false discovery rate using the command: FalseDiscoveryRate -in <merged.idXML> -out <fdr_out.idXML>.

The commands and parameters used for polypeptide identification were consistent with those described in Yang et al.’s study^11^.

### Identification of TST-expressed peptides in immunopeptidome data

In order to identify immunopeptides derived from tissue-specific transcripts (TSTs) using immunopeptidomic techniques, we retrieved TST protein sequences following the previously described method for neoantigen identification but without limiting their length to 9-mers. For HLA-pulldown sample analysis, MaxQuant version 1.6.3.4^60^ was used with a custom proteome database composed of the Uniprot reference database and TST protein-coding sequences. The search parameters were optimized for identifying TST-derived immunopeptides, including setting decoyMode to revert for reversing the peptide sequence as a decoy sequence, Carbamidomethyl (C) as the fixed modification, Acetyl (Protein N-term)/Oxidation (M) as variable modifications, and setting the minimum and maximum peptide lengths to 8 and 14, respectively, given that peptides that bind to MHC-I molecules are typically 8-14 amino acids long. To further maximize the identification of TST-derived immunopeptides, no digesting enzyme was specified. To verify that whether tsTE-derived immunopeptides were derived from Endogenous Viral Elements(EVE), we retrieved open reading frame (ORF) locations from EVEs in the gEVE database. A blastp analysis was performed between the gEVE protein sequences and the identified immunopeptidomic sequences to assess amino acid sequence similarity. No threshold on Evalue was set, and only tsTE-derived peptides with 100% similarity were retained.

### Correlation of TST-derived neoantigen burden and immunotherapy response

Seung Tae Kim et al. assessed the effectiveness of pembrolizumab in treating metastatic gastric cancer patients and categorized them into responders and non-responders to immunotherapy^61^. We obtained 45 RNA-seq raw data from metastatic gastric cancer under PRJEB25780 and used antigen prediction pipeline as described above to predict tumor neoantigens derived from TSJs/TSTs. ASJA was employed to identify splicing junctions of these samples, and the same criteria used for TST analysis for TCGA samples were applied to identify TSJs/TSTs which requires TSTs with frequency greater than 5% and are at least 10-fold higher expressed in tumor than normal or not expressed in normal samples. For each metastatic gastric cancer sample, we generated two individual polypeptide databases: TSJ database comprising MT peptide sequences with ten flanking amino acids on each side of the TSJs and TSTs database consisting of 9-mer coding protein sequences. Finally, netMHCpan-4.1 software was employed to predict TSJ/TST-derived neoantigens that could potentially bind to MHC-I molecules based on the HLA typing results obtained. To determine the gene expression levels of metastatic gastric cancer patients, FeatureCounts software^62^ was used to count the RNA-seq data, followed by normalization of the counts to transcripts per million (TPM) values using R programming language. Then the expression of immune checkpoint molecules was extracted for analysis.

### Identification and characterization of splicing junctions in extracellular vesicles

We detected and analyzed splicing sites in plasma EVs using exLR-Seq data through the following steps: (1) FastQC (version 0.11.8) was used to assess adapter removal quality and overall data quality. (2) Trimmomatic software^55^ (version 0.36) was utilized to remove Illumina adapter sequences and low-quality bases/reads. (3) ASJA was employed to identify and quantify linear splicing sites. HOMER was utilized to annotate the genomic locations of splice donor and acceptor sites, with the ‘bedtools intersect’ command used to define the host genes of the identified splicing sites. We observed that some splicing sites were present in EVs but not in tissues, while others were present in tissues but not in EVs. We performed KEGG pathway analysis on the source genes corresponding to these two groups of splicing sites using the ‘clusterProfiler’ R package^63^ to further clarify their functions.

### Identification of tissue and immune cell-specific splicing junctions

To investigate exosome sources, we identified tissue-specific and immune cell-specific splicing sites as signature junctions. To identify immune cell-specific splicing sites, we obtained raw RNA-seq data from 122 samples containing 30 types of immune cells from the Gene Expression Omnibus (accession number: GSE10701^64^) and utilized the ASJA pipeline to identify and quantify linear splicing sites in immune cell samples. We calculated the SPECS score for each splicing site based on its splicing expression matrix, and defined splicing sites with a SPECS score > 0.95 as immune cell-specific splicing sites. Furthermore, we identified tissue-specific splicing sites from the GTEx splicing profile using similar criteria (SPECS score > 0.95).

To trace the origin of exosomes, we filtered splicing sites using the following criteria to define signature junctions: (1) an average expression value > 1 in plasma EVs; (2) identification as immune cell/normal tissue-specific splicing junctions; and (3) for genes with multiple tissue-specific splicing sites, only the splice junction with the highest SPECS score was retained to avoid redundancy. We utilized the ssGSEA algorithm from the R package ‘GSVA’^65^ to compute immune cell/normal tissue enrichment scores for each plasma EV sample. The resulting scores were normalized to a range of 0-1 to indicate the proportion of EVs derived from each source.

### Classifier construction with RNA splicing junctions in EVs

To identify potential cancer biomarkers, cancer-specific splicing sites in Evs were identified for each type of cancer using the following criteria: Health_freq=0 & Cancer_freq>10%; Cancer_freq>5*Health_freq & Freq_cancer>10%. Here, Health_freq represents the expression frequency of a splicing site in EVs from healthy individuals, while Freq_cancer represents the expression frequency of a splicing site in EVs from cancer patients. Generally, splicing sites with high expression frequency in cancer and low expression frequency in healthy individuals were considered potential biomarkers. To distinguish different cancer types using splicing sites, we constructed SVM models based on RNA splicing levels in plasma EVs from 735 EV tumor samples. The samples were randomly divided into training and test cohorts at a ratio of 7:3. The specificity score were calculated of splicing across seven types of cancer using the ‘SPECS’ package in the training set and selected splicing sites with SPECS score >0.95 as the feature set. We generated an SVM model using the e1071 svm function with parameters type = “C”, kernel = “radial”, probability = TRUE. The performance of the model was assessed by calculating the area under the ROC curve (AUC) using the ‘pROC’ package in R. Finally, we selected the optimal model based on the maximum AUC value obtained from ten-fold cross-validation of the training and testing sets.

## Data availability

The ASJA program used can be accessed at https://github.com/HuangLab-Fudan/ASJA. The splicing sites and corresponding annotation information for cancer and normal tissues have been organized in RJunBase (http://www.rjunbase.org/). Data on TCGA samples, including TST score, stemness index, and other clinical information, are integrated and available in Supplement Table 1. Sample IDs for immune peptide profiling and identified immune peptides using MS-based immunepeptidome can be found in Supplementary Table 5. All other data supporting the findings of this study are available upon reasonable request from the corresponding author.

## Code availability

The bioinformatics analyses for this study were performed using various open-source software tools and packages. These include FastQC, STAR (v.2.5.2a), Trimmomatic (v.0.36), StringTie (v.1.3.4), FeatureCounts, FLAIR, 2passtools, deptools, SPECS, HOMER, clusterProfiler, CytoTRACE, Monocle, seq2HLA, NetMHCpan-4.1, OpenMS, MaxQuant, GSVA, e1071, pROC, circlize, survminer, ggplot2, samtools, ComplexHeatmap, and the Python package biopython. Additional processing involved in-house scripts, which are available upon request.

## Acknowledgments

This work was supported by grants from National Key Research and Development Project of China (2021YFA1300500) and National Natural Science Foundation of China (81872294, 82072694).

## Contributions

Conceptualization and writing-revision, S.H. and Z.H; Data curation, software, methodology, and analysis, Q.L.; Writing-draft, Q.L., Z.L.and H.L.; Visualization, Q.L. and J.Z; Experiments, Ziteng Li, H.Y., B.C. and Y.L.; Resources, H.Z.,J.H. and Z.M.; Funding acquisition supervision, and project administration, S.H.

## Conflict of intersts

The authors declare no competing interests.

## Supplementary figure legends

**Extended Data Fig 1. Tissue-specific splicing sites and gene expression of *NOL12*. a,** Tissue-specific splicing sites from *NOL12* and its parent gene expression (b).

**Extended Data Fig 2. Broad and specific expression of TSTs. a,** TST derived from XCL1 is widely expressed in various cancers. **b,** a TST derived from ID4 is specifically expressed in ovarian cancer.

**Extended Data Fig 3. Prognostic TSTs in each cancer type. a,** Number of TSTs associated with four clinical endpoints in each cancer type. **b,** TSTs can be associated with prognosis in multiple cancers. Rows are TSTs and columns are TCGA cancer types. Color indicates the total number of clinical endpoints associated with TST score, ranging from four (highest) to zero (lowest). Here we show the top 30 TSTs with the highest frequency. **c,** Association of TST score with four clinical endpoints in each cancer type. Red indicates association (p < 0.05); White indicates no association (p>0.05); Black indicates not evaluated due to insufficient number of samples (less than 5) with corresponding clinical information. **d,** At pan-cancer level, TST score was associated with DSS, PFI and DFI. The hazard ratios (HR) and p-values of logrank test were indicated at the top right corner.

**Extended Data Fig 4. Cancer types with significant correlation between TST score and stemness** (Pearson correlation coefficient >0.31, p < 0.05).

**Extended Data Fig 5. The characterization and oncogenic potential of tsTE1. a,** The expression of tsTE1 among pancancer tumor samples in TCGA. **b,** Relative subcellular localization of tsTE1 in MHCC-97L cells using U2 and β-actin transcript as nuclear and cytoplasmic control respectively. **c,** The amplification of tsTE1 for 5’ and 3’-RACE assays together with the full-length sequence of tsTE1. **d,** Cell colony counts estimation demonstrating the enhanced colony formation ability stimulated by stable overexpression of tsTE1 in various cancer cell lines. Data are shown as mean± SEM. Two-tailed student t-test was applied for statistical analysis. *, P < 0.05; **, P < 0.01; ***, P < 0.001.

**Extended Data Fig 6. Flowchart of neoantigen prediction from TSJ or TST sources.**

**Extended Data Fig 7. Affinity and abundance of tumor neoantigens generated by TSJ. a,** Affinity scores of TSJ-derived peptides and MHC-I molecules. **b,** Peptides were detectable in multiple samples, with the x-axis representing the number of tumor samples and the y-axis representing the expression frequency of peptides in the corresponding interval. **c,** The number of peptides derived from TSJ was positively correlated with the number of coding TSTs and negatively correlated with TMB. Each dot represents a TCGA tumor sample (**d**). **e,** Length distribution of annotated peptides in UniProt and peptides derived from TSTs. **f,** Sequence motif of annotated peptides (top) and peptides derived from TSTs (bottom).

**Extended Data Fig 8. TSJ/TST neoantigen load is associated with immunotherapy efficacy. a,** TSJ neoantigen load in responders and non-responders group. PD, SD, PR, CR represent disease progression, stable disease, partial response and complete response respectively. Neoantigen load is the number of predicted TSJ-derived immunopeptides. **b,** Expression of immune checkpoint molecules in high and low TSJ neoantigen load groups (using median as threshold). **c,** Number of responders and non-responders in different groups. HighTST is high TST neoantigen load group (above upper quartile), LowTST/ModTST is low/moderate TST neoantigen load group (below/between upper and lower quartiles). **d,** Difference in CD8+ T cell infiltration among different groups. HML represents high mutation load, HTST, MTST, LTST represent high/moderate/low TST neoantigen load.

**Extended Data Fig 9. Exploring splicing patterns and differential expression in cancer. a,** Enriched pathways of parent genes for splicing sites specifically detected in tumor tissue and EVs. **b,** Differential splicing between cancer and healthy samples. In the clustered heatmap, each row represents a single linear splicing site, and each column represents a sample. Expression values are normalized by row. **c,** Splicing sites with cancer-specific expression from tumor-associated genes. **d-f,** Splicing junctions with different frequency between liver, pancreatic, and breast cancer samples and control samples. The splicing junctions of interest were marked in red. Junctions were renamed for ease of reading according to the following rules: gene name + LS (linear splicing) + splicing sequencing ID (sorted by genomic location in genes).

**Supplementary Table 5:**
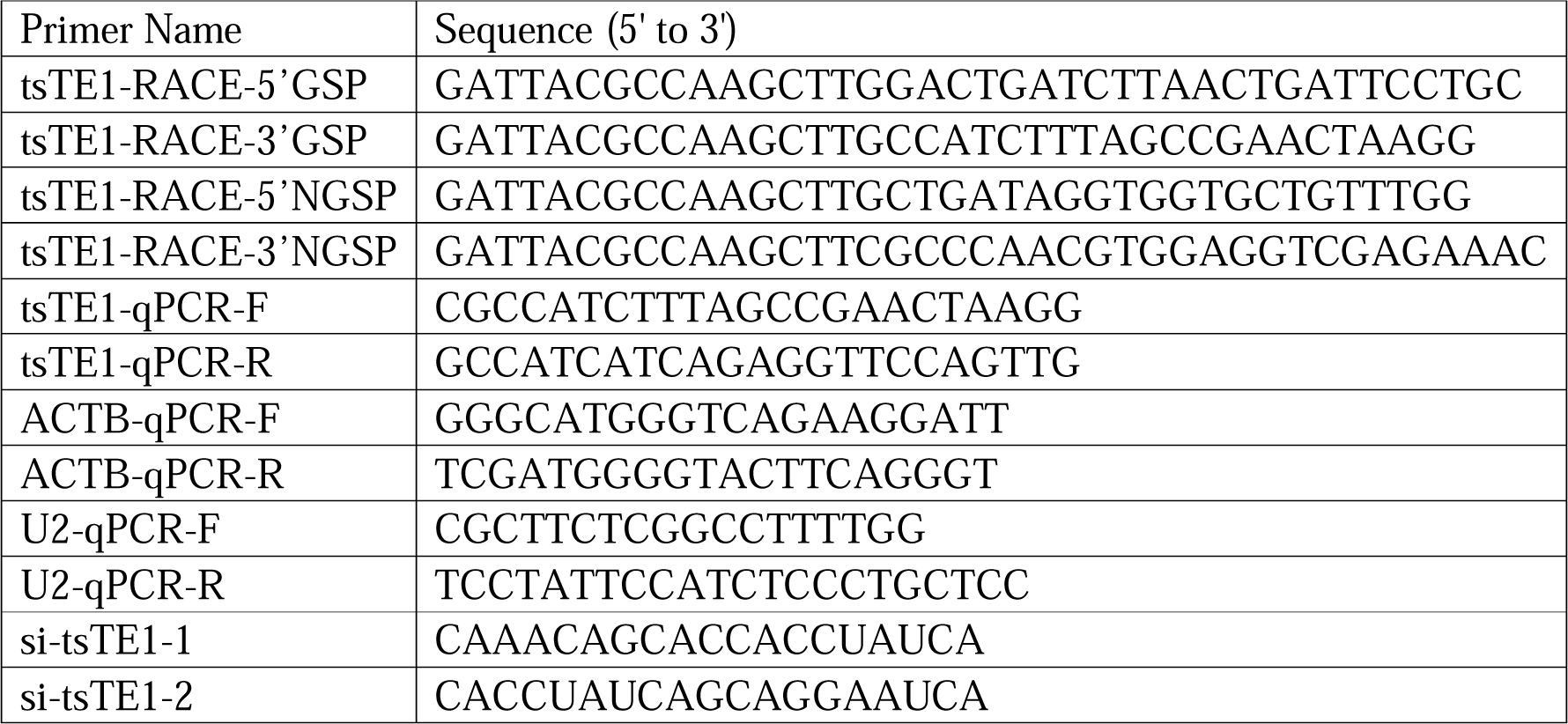
Sequences of primers and siRNA.

