## Supplementary figures and images for "RNA Splicing Junction Landscape Reveals Abundant Tumor-Specific Transcripts in Human Cancer"

### Extended data Fig1

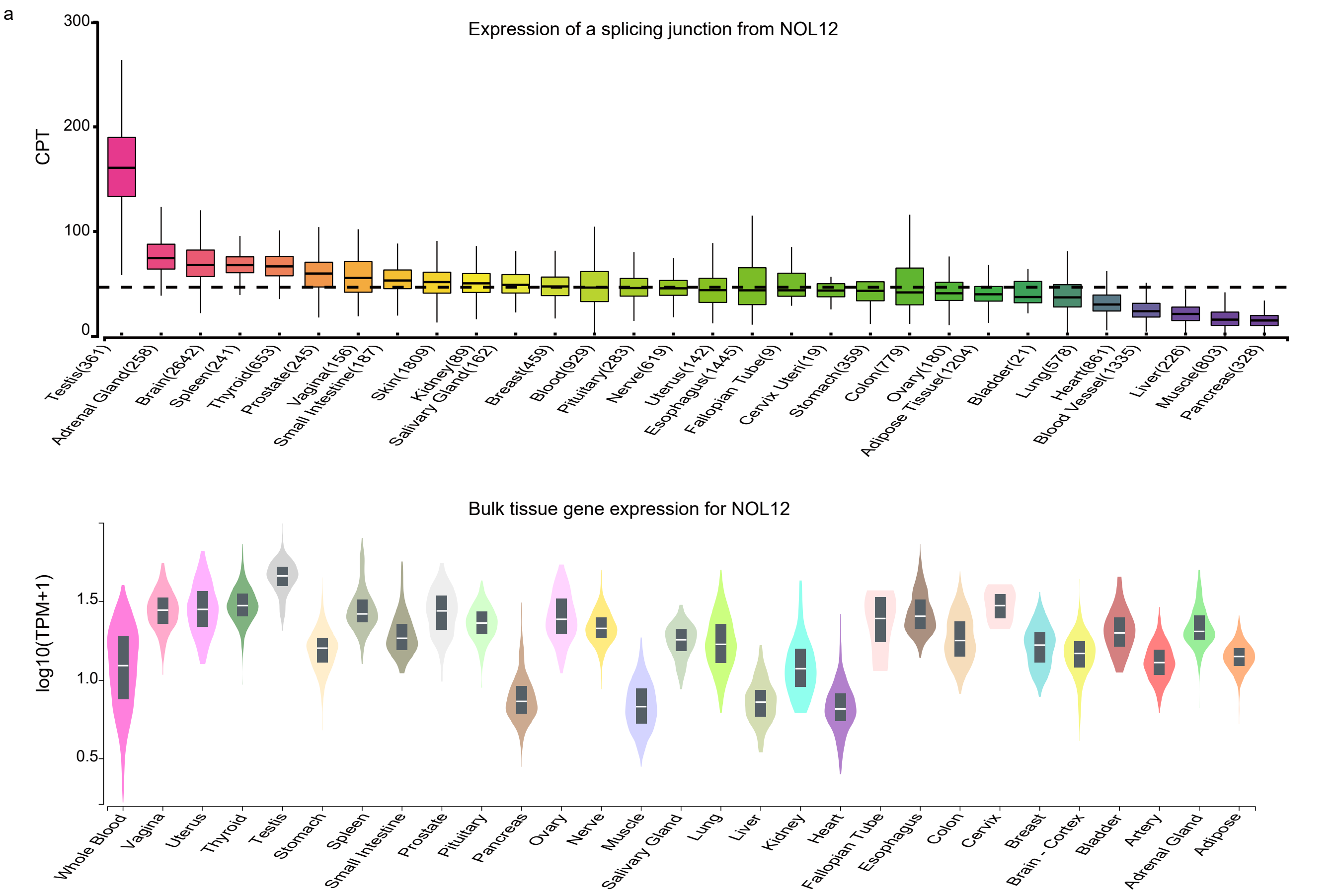

### Extended data Fig2

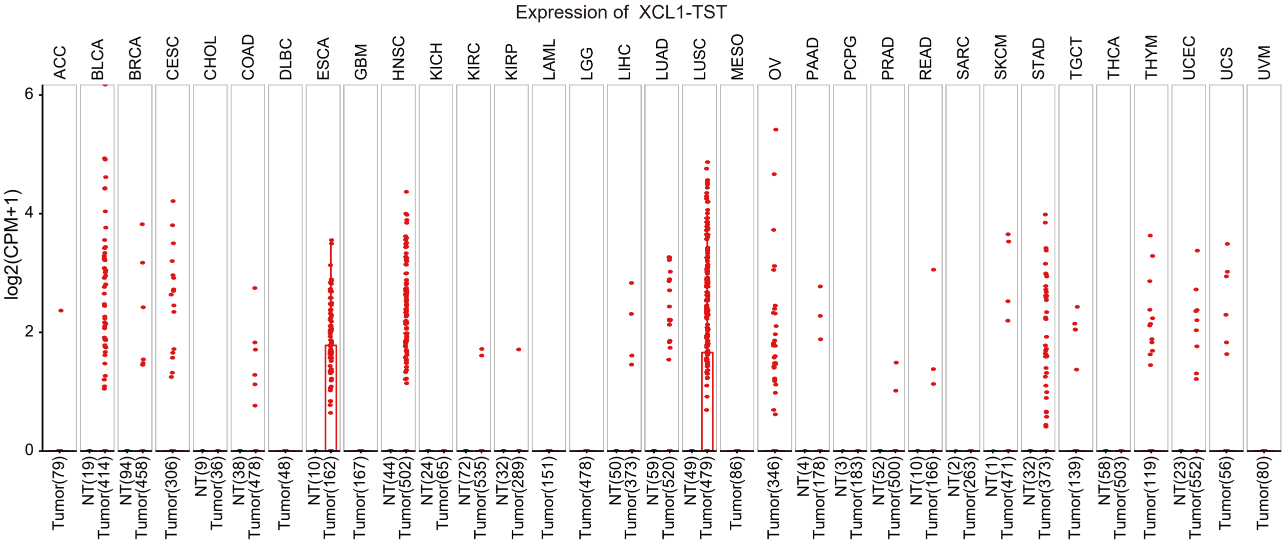

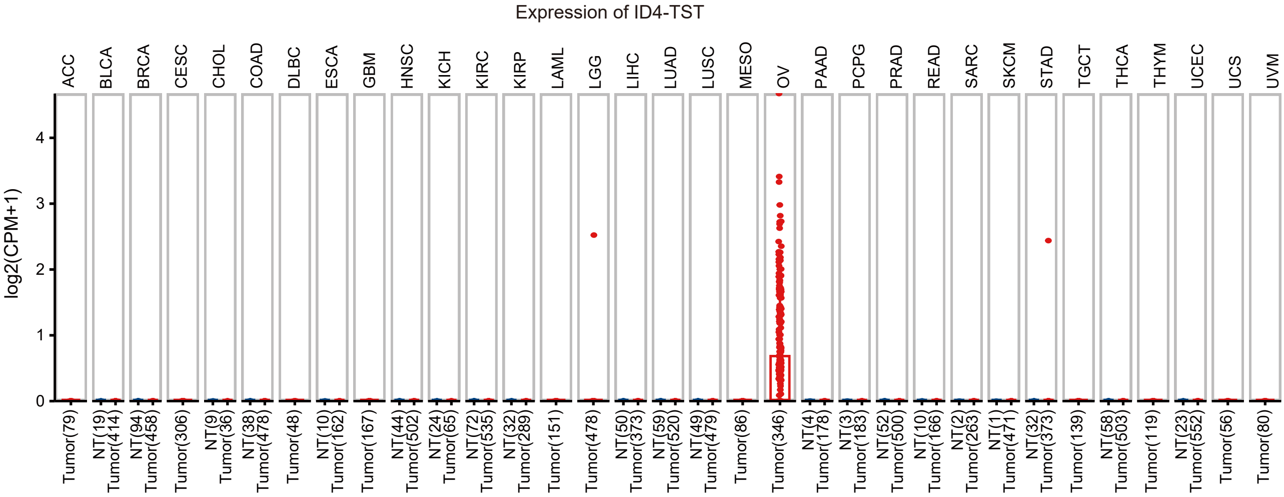

### Extended data Fig3

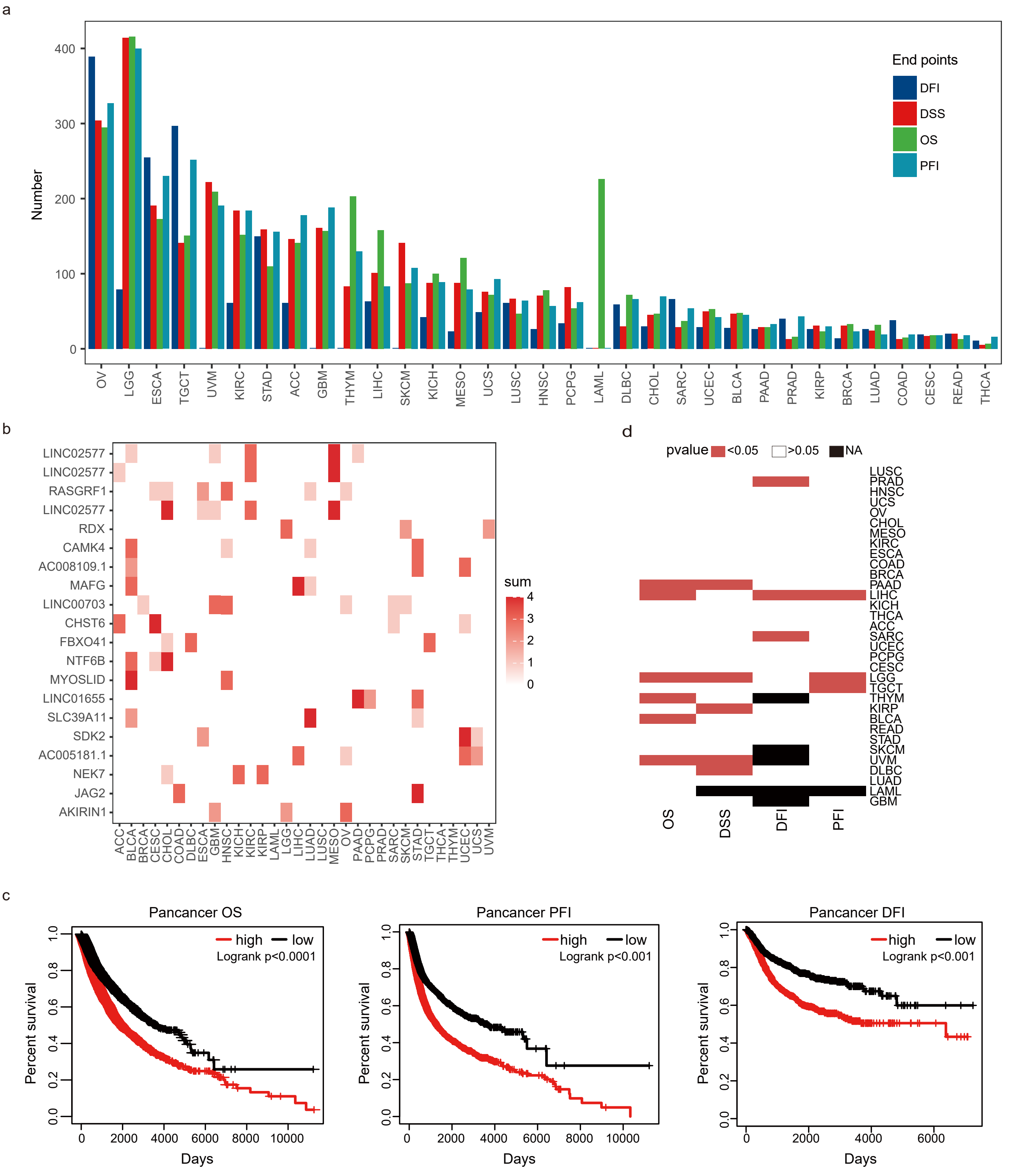

### Extended data Fig4

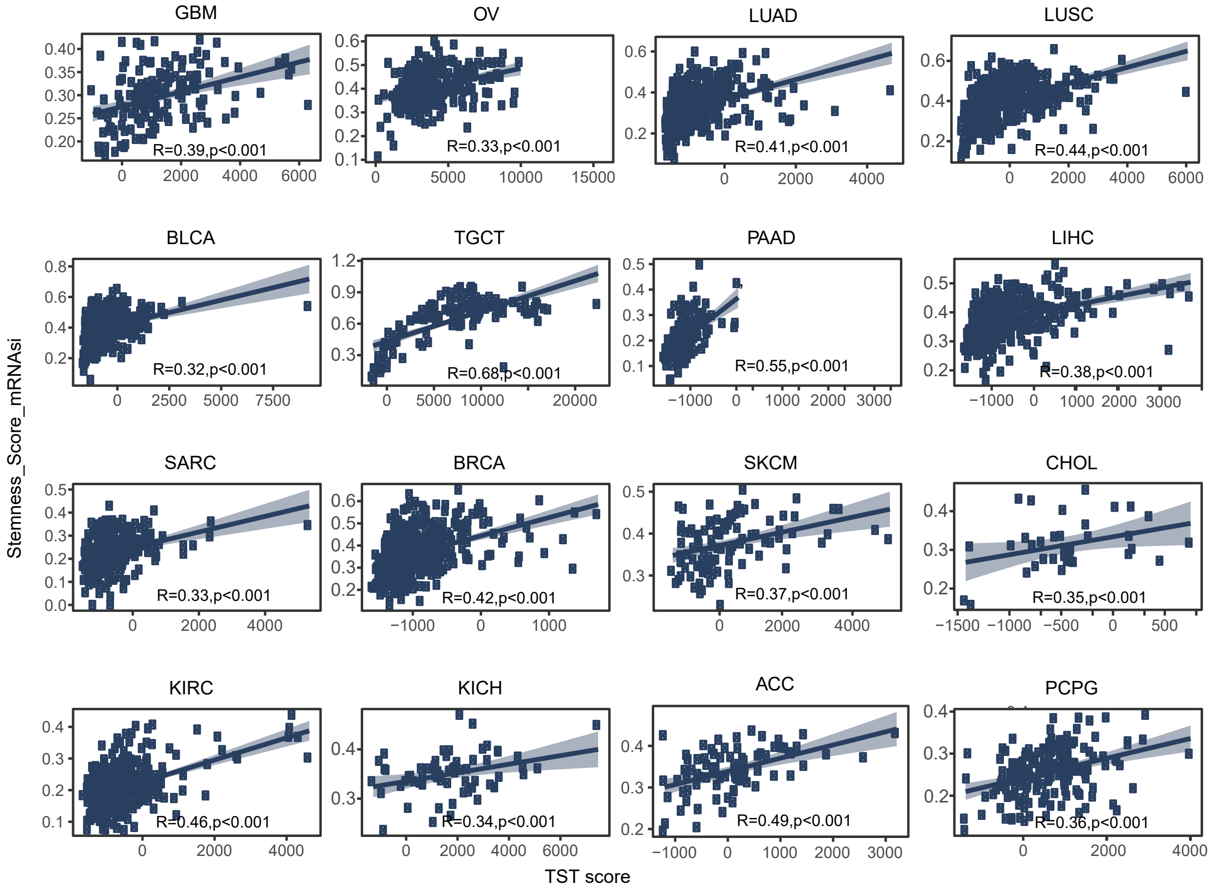

### Extended data Fig5

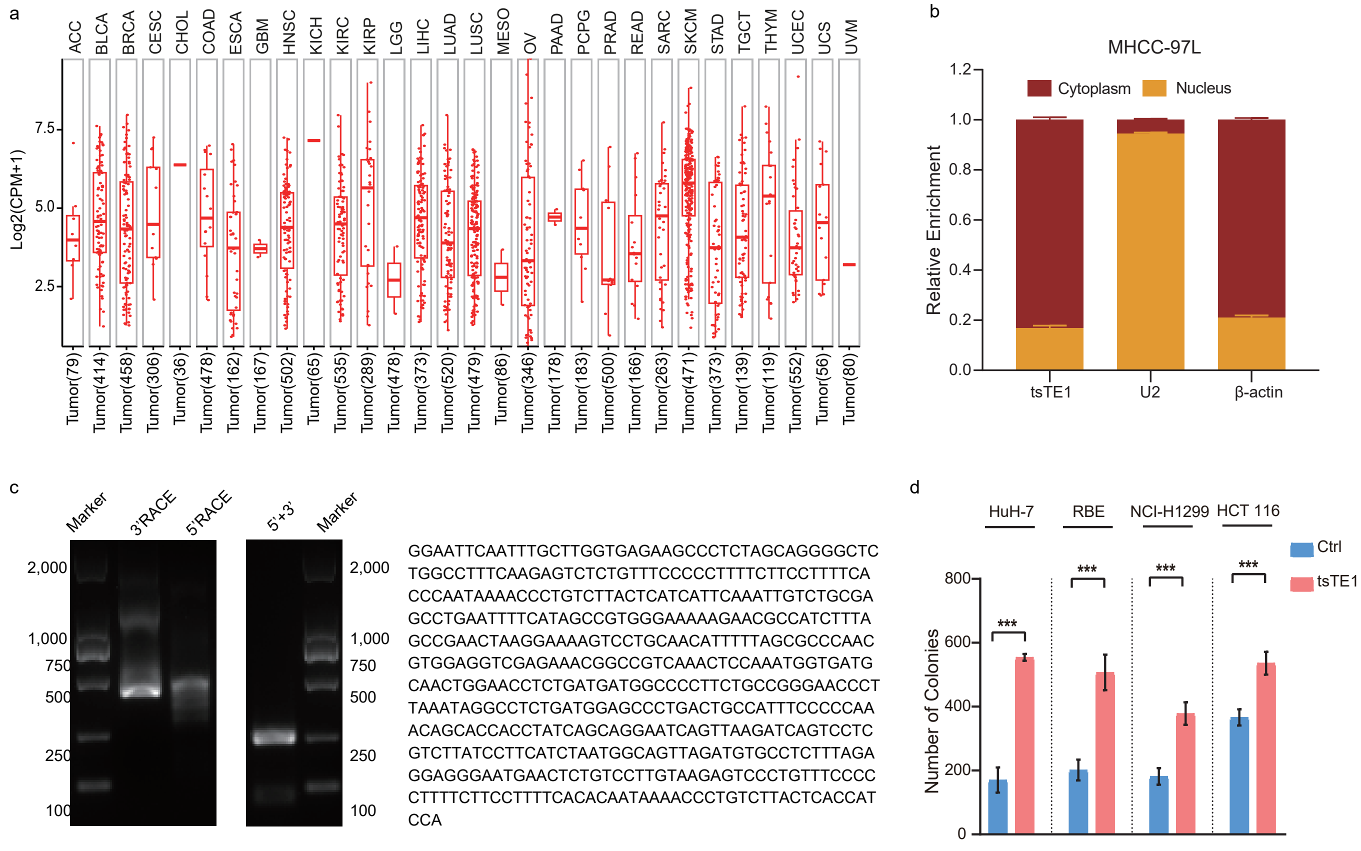

### Extended data Fig6

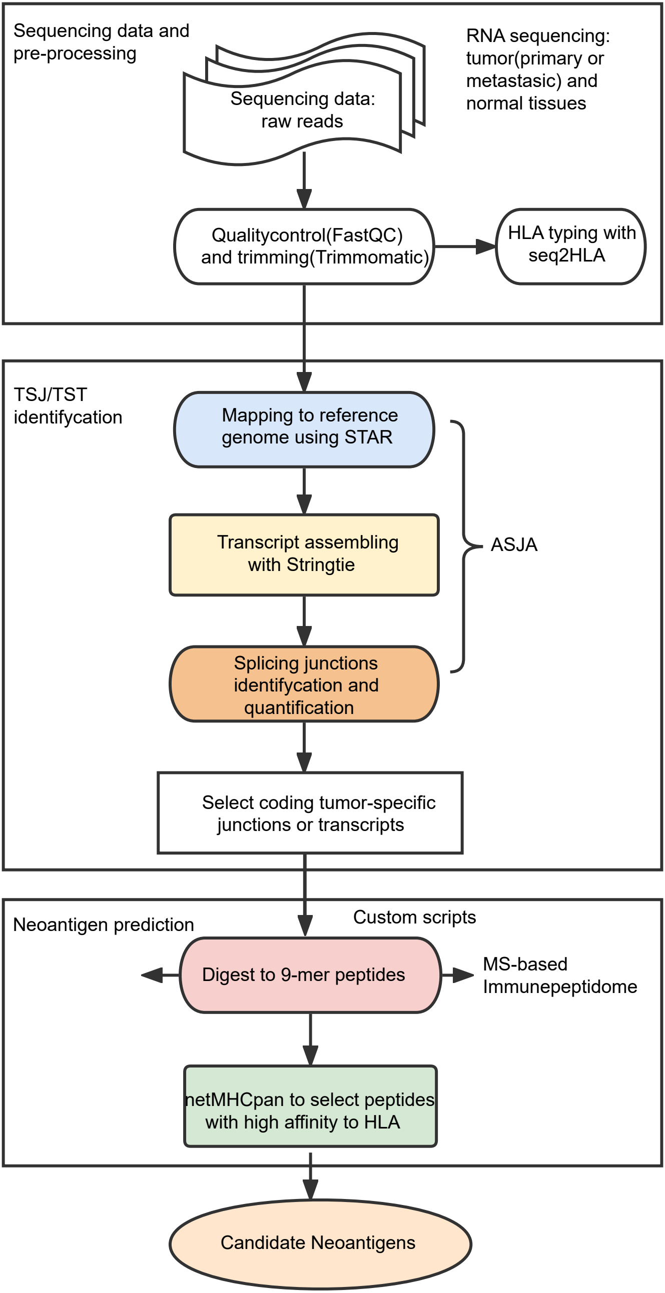

### Extended data Fig7

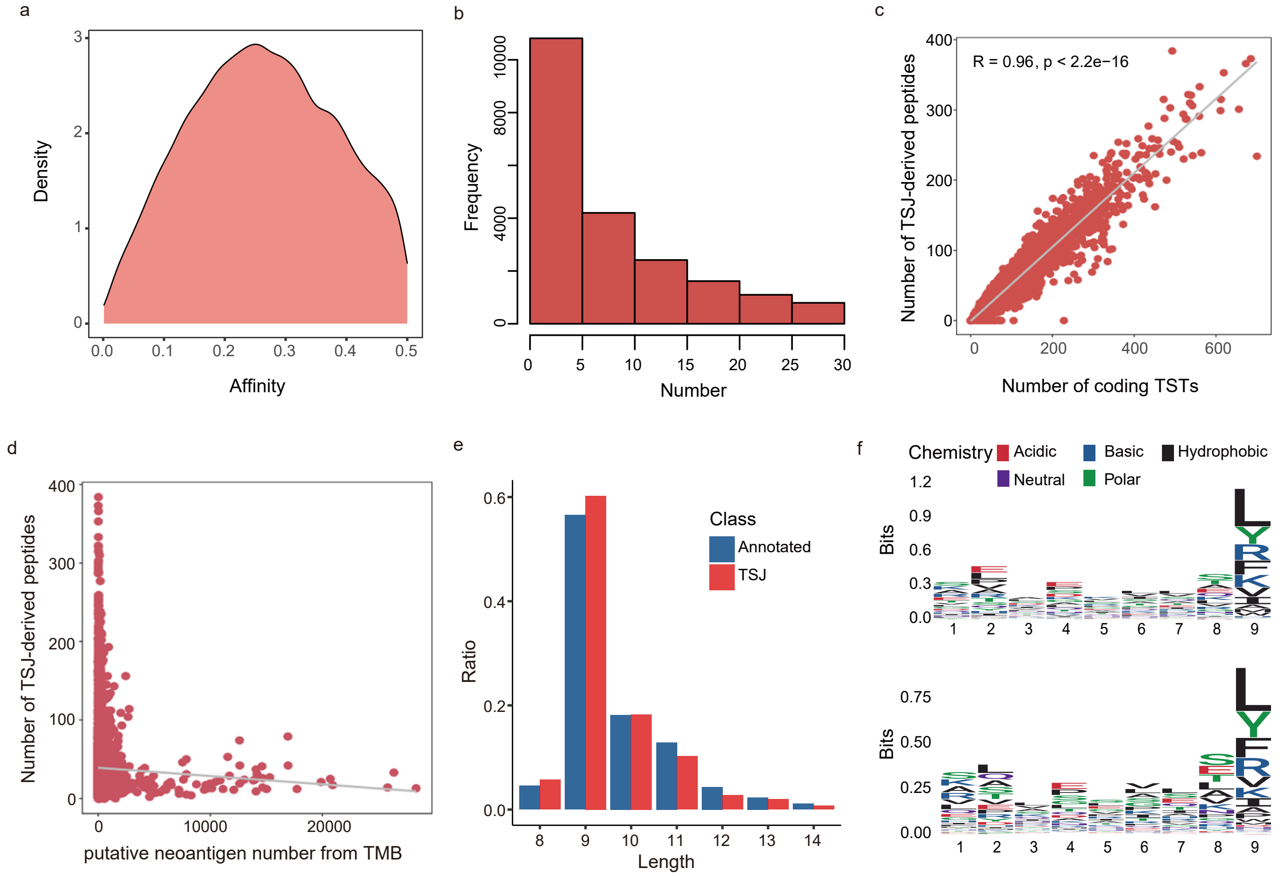

### Extended data Fig8

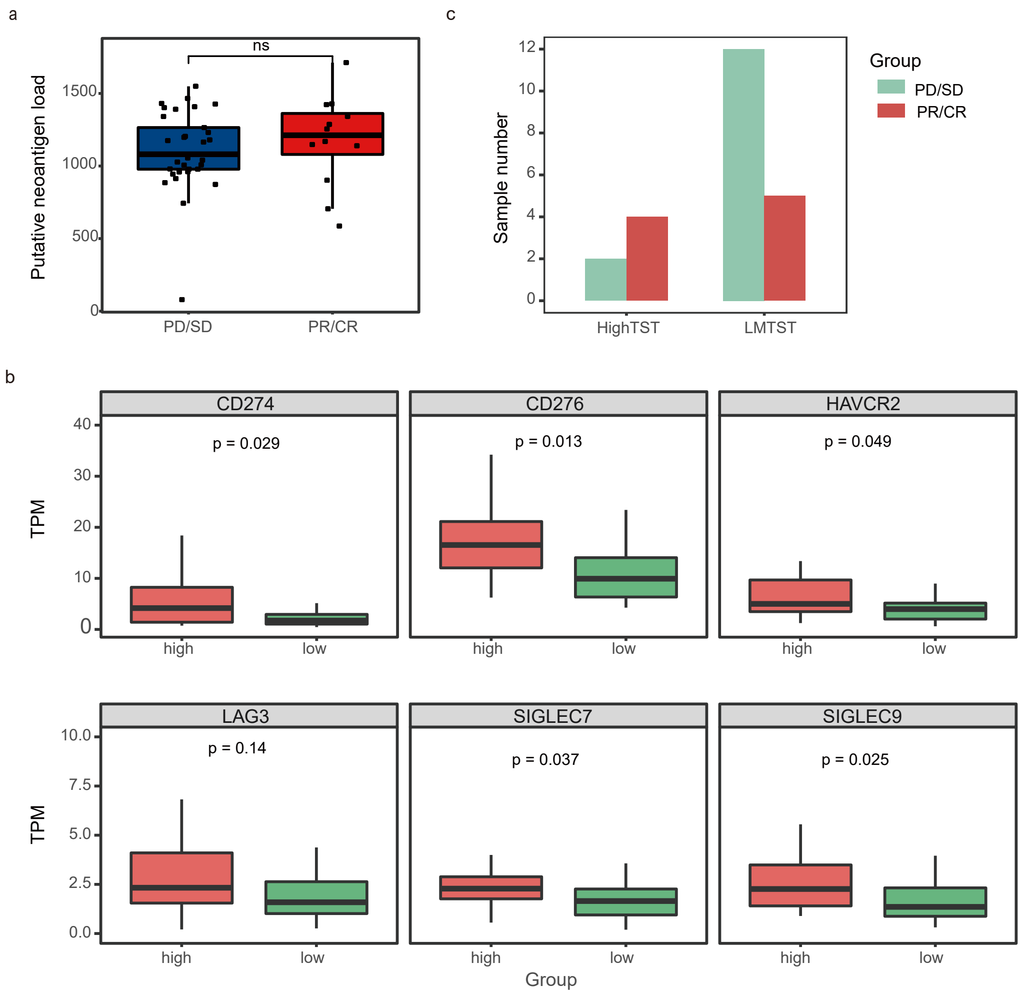

### Extended data Fig9

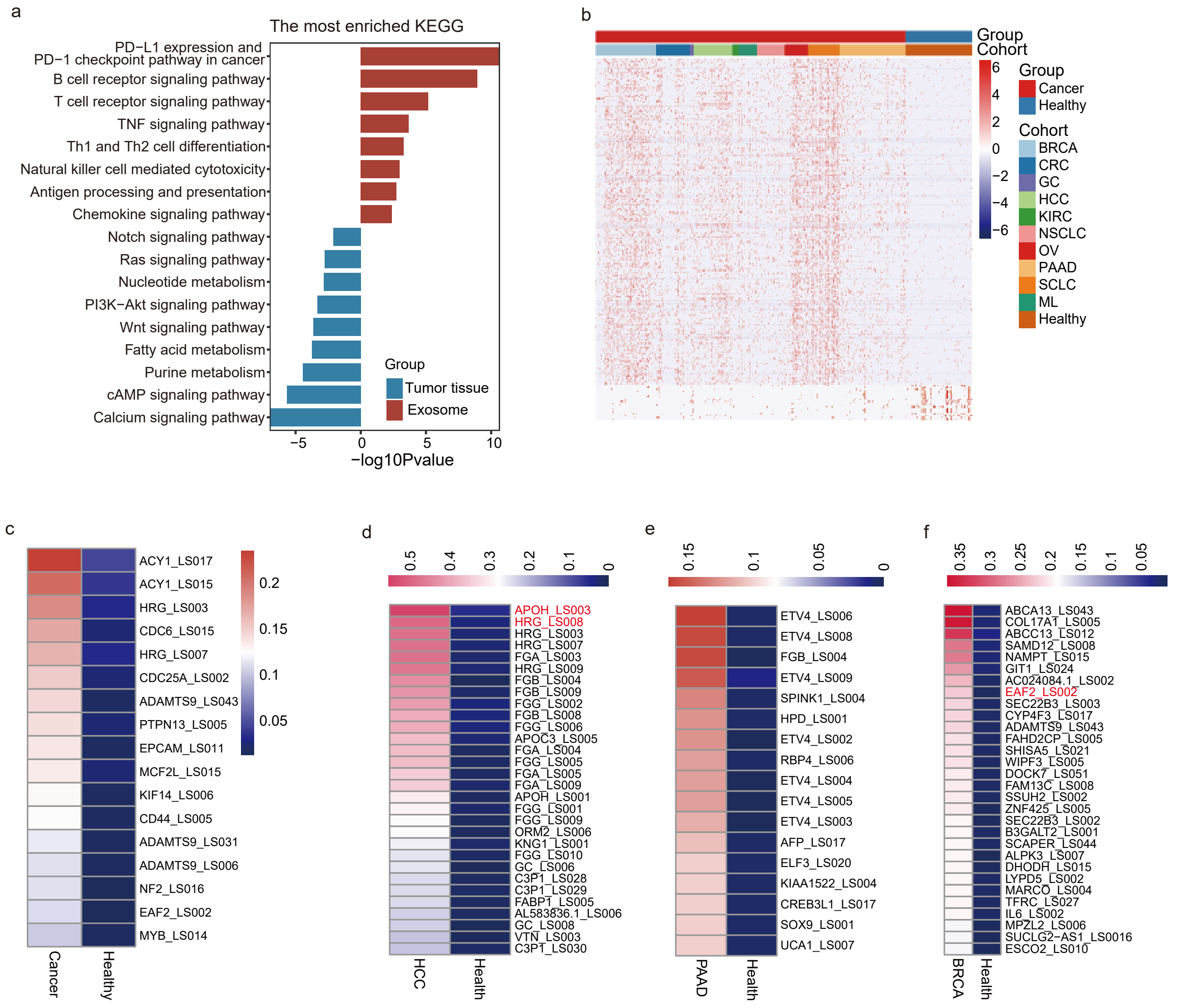
